# CLADES: a programmable sequence of reporters for lineage analysis

**DOI:** 10.1101/655308

**Authors:** Jorge Garcia-Marques, Ching-Po Yang, Isabel Espinosa-Medina, Minoru Koyama, Tzumin Lee

## Abstract

We present CLADES (Cell Lineage Access Driven by an Edition Sequence), a technology for cell lineage studies based on CRISPR/Cas9. CLADES relies on a system of genetic switches to activate and inactivate reporter genes in a pre-determined order. Targeting CLADES to progenitor cells allows the progeny to inherit a sequential cascade of reporters, coupling birth order with reporter expression. This gives us temporal resolution of lineage development that can be used to deconstruct an extended cell lineage by tracking the reporters expressed in the progeny. When targeted to the germ line, the same cascade progresses across animal generations, marking each generation with the corresponding combination of reporters. CLADES thus offers an innovative strategy for making programmable cascades of genes that can be used for genetic manipulation or to record serial biological events.

**One Sentence Summary:** A sequence of reporters for lineage analysis

## Main Text

Cell lineage is an essential determinant in the acquisition of cell identity (1–2). Establishing the association between cell lineage and fate is one of the fundamental challenges in biology. Solving this puzzle will provide a unique framework to interrogate the molecular mechanisms involved in cell type specification: it is not possible to fully understand how a molecular factor affects cell fate decisions if we do not even know where/when these cell fate decisions occur.

While single-cell transcriptomics has made it possible to identify cell types with great detail, tracing lineages in complex organisms remains challenging. Two main strategies have been deployed: i) imaging-based methods that label cell lineages in intact tissues (3–4), and ii) DNA sequencing-based methods which unravel cell lineages via phylogenetic analysis of DNA mutations accumulated during development (5–7). Unless the specimen is accessible for real-time visualization, imaging-based strategies are only able to label cell clones rather than tracing lineage progression in a single individual. For organisms with stereotyped development, full lineages can be assembled by resolving smaller segments in multiple individuals (8). Besides overlooking inter-individual differences, this approach is tedious and makes it impractical to analyze mutant lineages in detail. On the other hand, methods based on DNA sequencing allow high-throughput analysis of lineage progression, although the resolution is currently limited to major lineage branches (7). Moreover, sequencing methods fail to recover spatial and morphological information, critical features for identification of cell types and mutant phenotypes.

To circumvent these limitations, we developed CLADES, a technology based on CRISPR/Cas9 to trace and manipulate lineages in *Drosophila*. Inspired by principles of synthetic biology, we engineered a programmable system of genetic switches to control the activation and inactivation of reporter genes in a coordinated manner. This creates a sequence of colors (reporter combinations) whose order is pre-established. Birth-order is then deducted from the specific color inherited by the progeny (neurons) from the progenitor cell (neuroblast). Thus, early- and late-born cells are labeled with the first or the last color in the sequence respectively. The coordinated activation and inactivation of reporters maximizes the number of markers that can be simultaneously imaged, permitting straightforward lineage tracing.

### A conditional gRNA scaffold that is activated by other gRNAs

Controlling the sequential activation and deactivation of multiple reporter genes requires a sophisticated mechanism of genetic switches, unattainable with any existent technology. We therefore decided to build upon CaSSA, a tool to control the activation of reporter genes (9). This technology utilizes CRISPR/Cas9 (10) to induce a double-strand break (DSB). If there are homologous sequences (direct repeats) on either side of the DSB, single strand annealing (SSA) is the most likely mechanism of DNA repair (11–12). This pathway repairs DNA by recombining both direct repeats and removing the intervening sequence. As this repair outcome is predictable, one can design genetic elements that only become active after a DSB is repaired by SSA. Despite we initially applied it to reporter genes (9), we reasoned this concept could be extended to generate gRNAs that could be controlled by other gRNAs. These conditional gRNAs could then be organized as a synthetic cascade to sustain the expression of specific combinations of genetic markers in a pre-defined order.

To generate such conditional gRNAs, we need to make an inactive form of gRNA with two direct repeats flanking the target site of a trigger gRNA (switch). In the initial, inactive gRNA configuration, the inclusion of the switch alters the secondary structure, thus abolishing its activity. We first explored various strategies to make such inactive form of gRNA (Fig. S1), resulting in an optimized conditional gRNA containing the switch within the scaffold region (Fig. 1A).

**Fig 1.**
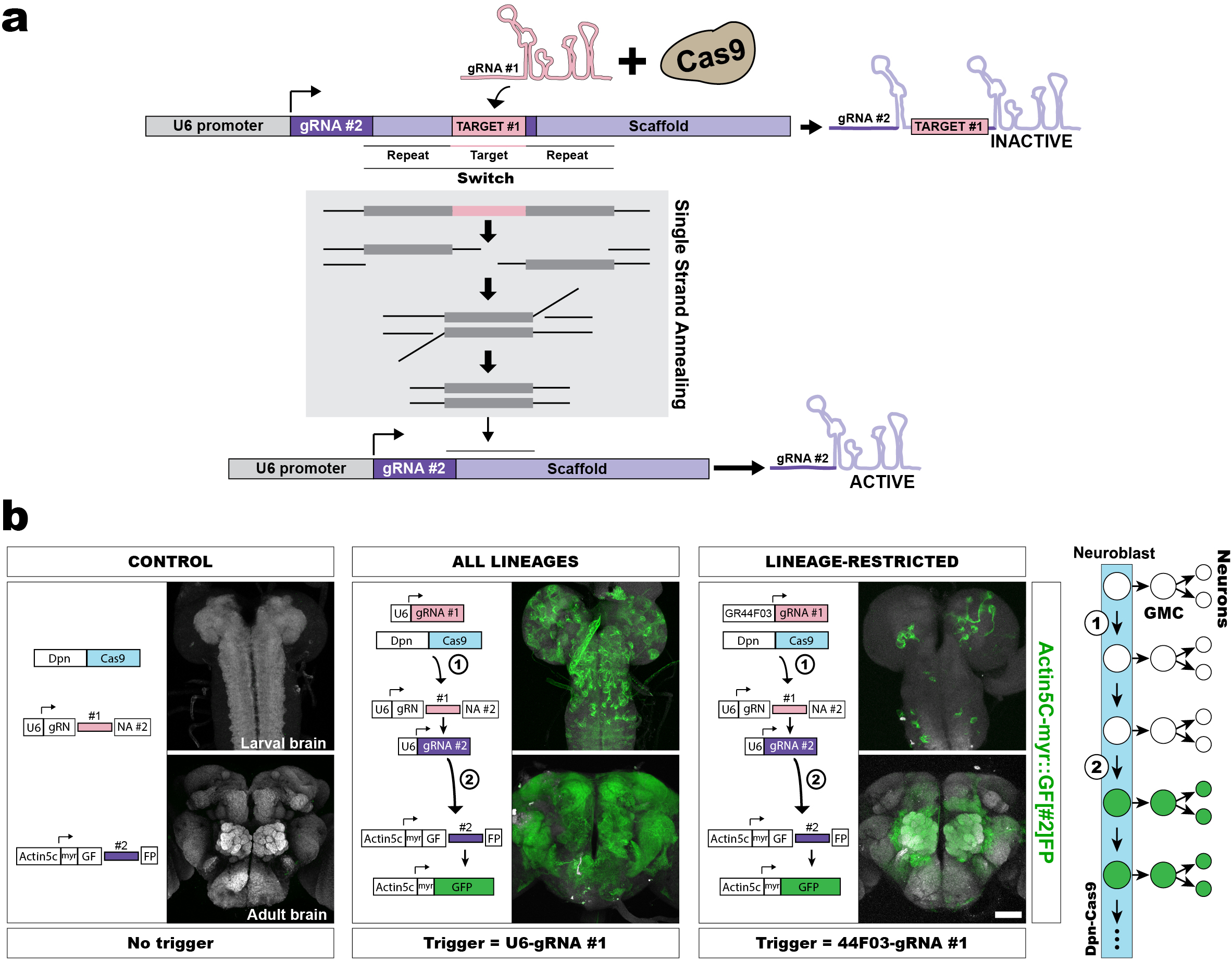
A conditional gRNA scaffold that is activated by other gRNAs. (A) Cartoon illustrating the conditional gRNA design. (B) Conditional gRNA activation by a trigger gRNA. For each experiment: left, scheme showing the cascade of events occurring in the fly brain. Right, representative examples of larval and adult brains showing immunohistochemistry for GFP expression. N=30 brains. Scale bar = 50 micrometers.

The switch is initiated only after the induction of a DSB by the combination of Cas9 and a trigger gRNA. The ensuing SSA repair recombines both repeats and collapses the switch, restoring the original gRNA structure and function. To validate the optimized conditional gRNA (Fig. 1B), we generated a fly line expressing three transgenes: i) the conditional gRNA, gRN[#1]NA#2 under the control of the ubiquitous U6 promoter. The nomenclature of the conditional gRNAs reflects the target site for the specific trigger gRNA within brackets and the direct repeats surrounding the target are represented by a repeated letter, ii) a conditional GFP reporter that responds to the gRNA#2 (9), GF[#2]FP driven by ubiquitous Actin5C, and iii) a Cas9 nuclease driven by a neuroblast-specific promoter, deadpan (Dpn). If, as we envision, the conditional gRN[#1]NA#2 is inactive in the initial configuration, the GF[#2]FP reporter will also remain inactive in the absence of the trigger gRNA. We found only minimal leakiness when the conditional gRNA and conditional GFP were co-expressed without the trigger gRNA (∼1 out of 4.72×10^4^ neuroblasts had GFP fluorescence). Only after crossing the line above to a fly bearing the trigger U6-gRNA#1, the conditional gRN[#1]NA#2 became functional and, in turn, activated the GF[#2]FP reporter, resulting in abundant GFP expression. Given that the reconstitution of the reporter is an inheritable modification in the DNA, confining Cas9 to the neuroblasts resulted in GFP expression in these cells but also in their progeny. This led to a seemingly ubiquitous expression of GFP in adult brains. We also confirmed that this expression could be restricted to a subset of lineages by providing the trigger gRNA#1 under the regulation of GR44F03, a driver that is mostly restricted to the antennal lobe lineages (9). This resulted in the expected pattern with consistent labeling of antennal lobe lineages plus occasional hits in mushroom body and ellipsoid body lineages (Fig. 1B). To directly quantify the extent of gRNA reconstitution, we examined the gRNA repair outcome by targeted amplicon analysis via next-generation sequencing (Fig. S2). The adult brains retained a minor proportion (14.17%) of the conditional gRN[#1]NA#2 in the unedited state, and SSA repair occurred in 56.28% of the edited reads. The rest of edits consisted mostly of small deletions or insertions. Despite the suboptimal efficiency, we reasoned this would not be an issue as CLADES should allow to resolve a lineage from a small number of clones (see below). These results confirm the success of the conditional gRNA design, which could respond to other gRNAs with minimal leakiness. To explore the applicability of this technology to vertebrate models, we tested the same conditional gRNA design in zebrafish (Fig. S3). We injected plasmids encoding: i) an injection control (RFP), Cas9 and a YF[#2]FP reporter, all of them in the same open reading frame transcribed from the Ubi promoter (ubiquitous) and ii) the U6-gRN[#1]NA#2 conditional gRNA. Only after providing the trigger gRNA#1 (Fig. S3B), a substantial percentage of cells expressed YFP (∼54%). This experiment proved that our conditional gRNA design worked consistently in fish, demonstrating the feasibility of this technology in vertebrates.

### A coupled cascade of gRNAs and reporters

Tracing lineages requires labeling cells with as many distinguishable marks as possible. For imaging-based methods, this parameter can be maximized via the specific combination of different fluorophores. The new conditional gRNA unlocks the potential to trigger cascades of gRNAs, which could in turn control the sequential activation/inactivation of reporter genes. A requisite to deliver a fixed sequence of reporter expression is that the order of reporter activation/inactivation should always follow the same order as the gRNA cascade. However, when the activation of reporters was controlled in trans by a gRNA cascade, we found this was not the case (Fig. S4). To circumvent this issue, we optimized a construct where the activation of conditional gRNAs was inextricably linked to the activation/inactivation of a reporter gene (Fig. 2 and S5). We accomplished this by embedding two conditional gRNAs within the open reading frame (ORF) of the reporter (Fig. S5), each comprising either the ON or the OFF cassette and controlling the activation or the inactivation of the reporter. In the initial state of the ON cassette, the switch in the conditional gRNA simultaneously inhibits the gRNA function and places the reporter out of frame. Coexistence of Cas9 and the trigger gRNA produces a DSB in this switch followed by SSA. The SSA event concurrently reconstitutes the conditional gRNA and brings the reporter into frame. Likewise, a similar switch in the OFF cassette controls the simultaneous activation of another gRNA and inactivation of the reporter. Based on this design, we generated a fly line expressing this conditional version of an RFP reporter (CLADES1.0-RFP). We also generated control flies expressing CLADES1.0-RFP variants with either the ON or both ON and OFF cassettes artificially pre-activated. As expected, the CLADES1.0-RFP-ON variant showed strong fluorescence and the CLADES1.0-RFP-ON-OFF variant had no fluorescence (Fig. S6). To test activation by the ON cassette, the line expressing CLADES1.0-RFP was crossed to flies bearing Dpn-Cas9 and the trigger U6-gRNA#1 (Fig. 2A). Only those flies expressing the three elements exhibited red fluorescence. This fluorescence was strong and seemingly ubiquitous in the brain. To test the OFF cassette, we crossed CLADES1.0-RFP-ON to a line with Dpn-Cas9 and the U6-gRNA#3, which eliminated most of the fluorescence in the fly brain (Fig. 2B). The residual fluorescence was expected and was most likely the consequence of: i) actin driven reporter expression in cells not derived from neuroblasts, ii) the editing event occurring after the birth of some early neurons, or iii) the occurrence of imprecise DNA repair that does not change the reporter ORF. These results prove that specific gRNAs can activate or inactivate a CLADES-reporter in an efficient manner.

**Fig 2.**
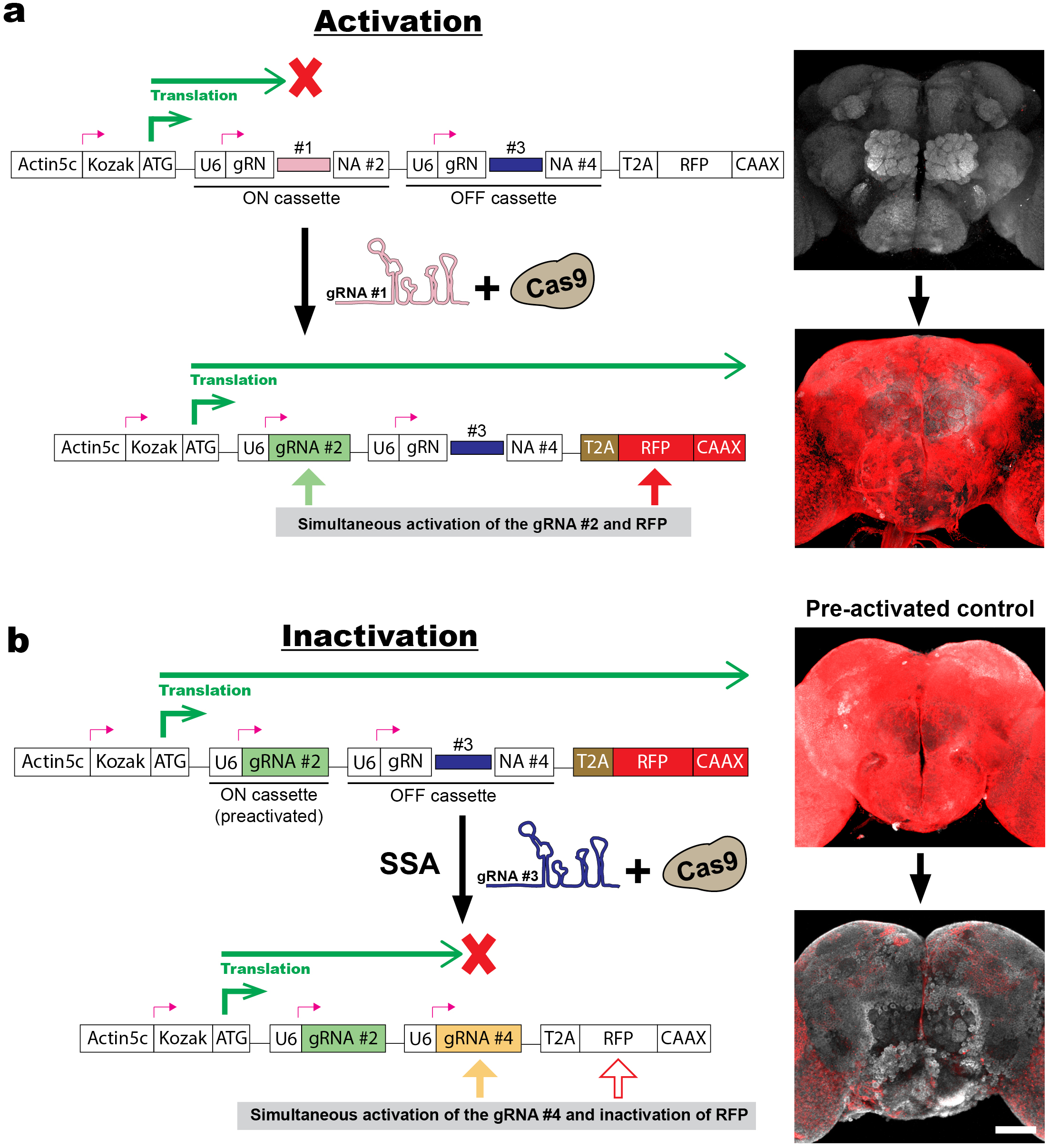
A coupled cascade of gRNAs and reporters. (A) The activation of the conditional gRNA#2 (ON cassette) by Cas9 and the trigger gRNA#1 leads to an ORF shift that brings the reporter into frame. Thus, both the gRNA#2 and the reporter (red fluorescence) become simultaneously active. (B) In the inactivation (or activation of the OFF cassette), the gRNA#4 is activated by Cas9 and the gRNA#3, bringing the reporter out of frame (tested with a pre-activated version of the construct). Red, immunohistochemistry for RFP. N=18 brains. Scale bar = 50 micrometers.

### CLADES 1.0: two reporters, three colors

We next examined if those conditional gRNAs, embedded within the ORF of reporter genes, can progress as a cascade. We designed the first reporter cascade based on CLADES (Fig. 3A). To do this, we generated a second reporter, CLADES1.0-YFP that would function together with CLADES1.0-RFP to sequentially change colors from green to yellow to red. In CLADES1.0-YFP, the gRNA#1 (gRNAs were renumbered for simplicity, see Table 2) is designed to concurrently turn on a YFP gene (YFP+ = green fluorescence) and the gRNA#2. In the subsequent step of the cascade, the gRNA#2 would trigger the ON cassette in CLADES1.0-RFP (YFP+/RFP+ = yellow fluorescence) as well as the gRNA#3. In the final cascade step, the gRNA#3 is intended to simultaneously activate the gRNA#4 and inactivate the YFP reporter (YFP-/RFP+ = red fluorescence). To test this cascade, we generated a fly bearing CLADES1.0-YFP, CLADES1.0-RFP, Dpn-Cas9 and U6-gRNA#1 transgenes. Neither the U6-gRNA#1 nor the Dpn-Cas9 transgene alone was able to trigger either reporter (Fig. S7A). Only after combining all four components (CLADES constructs are inserted into a single chromosome to facilitate future experiments), we found clones expressing green, yellow and/or red fluorescence. Similarly, in the absence of the first reporter, providing the U6-gRNA#1 and Dpn-Cas9 failed to activate the second reporter (Fig. S7B). We followed the color progression across larval development (Fig. S8). We detected the virtually only-green neuroblasts at 0H ALH and the plateauing of yellow neuroblasts at 24H ALH. The onset of RFP expression was followed by emergence of red-only neuroblasts at 60H ALH. The percentage of red-only neuroblasts increased at 84H ALH, concomitant with a decrease in yellow neuroblasts after the green reporter becomes inactive. The proportion of cells with each color decayed following the cascade progression, consistent with unedited reporters or expected indels that prevent the cascade from progressing to the next step.

**Fig 3.**
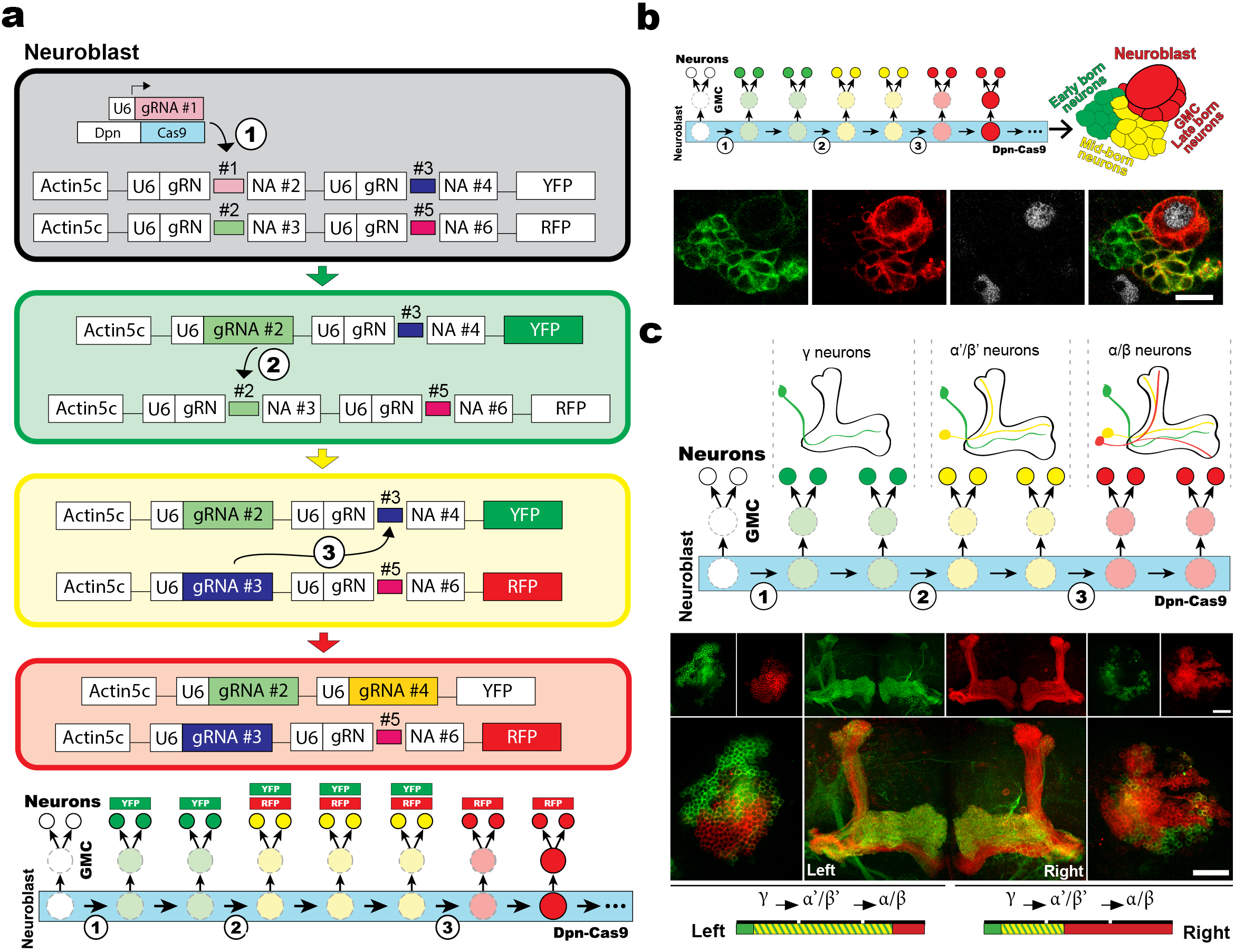
CLADES 1.0: two reporters, three colors. (A) Scheme illustrating the cascade progression in CLADES 1.0. (B) Neuronal clone in the larval brain labeled with CLADES. (C) CLADES labeling in the mushroom body lineage. In the right hemisphere, the red-only fluorescence labels the full alpha/beta and part of the alpha’/beta’ population, while the green and red or green-only labels the gamma and part of the alpha’/beta’ population. Green, red, gray, immunohistochemistry for GFP (YFP), RFP, Dpn respectively. Scale bars = 10 micrometers in B and 50 micrometers in C.

Now that we have a three-color cascade, we wanted to apply it to follow the progression of neuronal lineages. For an initial characterization, we tested CLADES using the GR44F03 driver so that we could resolve single lineages. In third instar larval brains, a typical pattern existed of neuroblasts expressing red fluorescence, surrounded by its progeny of GMCs and neurons labeled in red, yellow or green (Fig. 3B). Given minimal neuron migration in *Drosophila*, neurons are pushed away from the neuroblast as new GMCs and neurons are born. Consequently, closest to the neuroblast are the youngest cells, newborn GMCs and neurons (red), slightly further away are mid-born neurons (yellow) and finally the oldest, early-born neurons (green). To further characterize CLADES 1.0, we targeted it to the four ‘equivalent’ mushroom body lineages (Fig. 3C). These lineages are well characterized to give rise to three types of neurons in a consecutive order: gamma, alpha-prime/beta-prime, and finally alpha/beta neurons (13). For stochastic labeling, we used the minimal activity of the GR44F03 driver in the mushroom bodies to target only one of these lineages at a time. Notably, 3-color CLADES revealed a pattern consistent with the birth order previously described (Fig. 3C). In summary, these results demonstrate that gRNAs embedded in the CLADES constructs are functional and allow to design genetic cascades. Targeting CLADES to progenitor cells allowed the progeny to inherit a sequential cascade of reporters, coupling birth order with reporter expression.

### CLADES 2.0: combining a five-color cascade with GAL4 induction

Resolving cell lineages without cell type information provides limited insight. For systematic characterization and targeting of cell types, the fly community has exploited the GAL4/UAS system extensively (14). Combining CLADES with GAL4/UAS would allow concurrent interrogation of lineage progression and cell identity. We therefore constructed a new version of CLADES (2.0) compatible with the GAL4/UAS system. In order to cover the cells from the beginning of the lineage, CLADES 2.0 includes a pre-activated first reporter (CLADES2.0-YFP-ON) which can trigger the cascade of subsequent reporters in all progenitor cells. Moreover, we sought to extend the traceability by adding a third reporter (CFP), creating a 4-step cascade that could produce five colors in order: green, yellow, red, purple, blue (Fig. 4A). Importantly, these reporters are under UAS control and thus can be expressed only in GAL4-positive cells, although GAL4 is not required for the cascade progression. To validate CLADES 2.0 in all the neuroblast population, we crossed a fly bearing Dpn-Cas9 and the three CLADES constructs (CLADES2.0-YFP-ON, CLADES2.0-RFP, and CLADES2.0-CFP) to flies with Dpn-GAL4 (Fig. 4B, S9).

**Fig 4.**
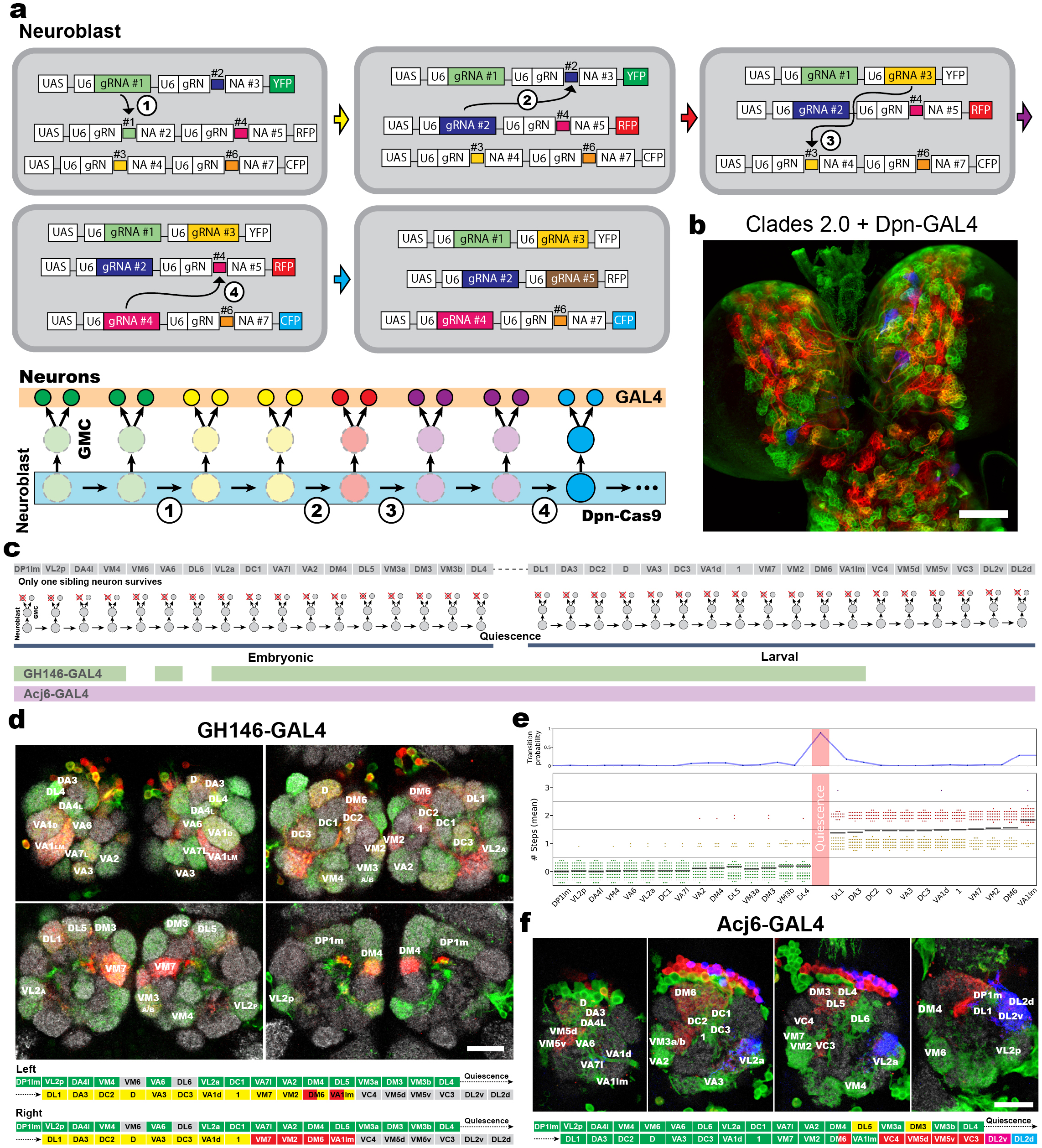
CLADES 2.0: combining a five-color cascade with GAL4 induction. (A) Cartoon illustrating the principle of CLADES 2.0. (B) Larval brain (wandering larva/white pupa) showing all neuroblasts labeled by combining CLADES 2.0 with Dpn-GAL4. (C) Scheme showing the order of neuronal types generation in the ALad1 lineage. In this lineage only one of the neurons arising from the GMC survives. Whereas Acj6-GAL4 can label the entire lineage, GH146-GAL4 labels only an early window. (D) Representative examples of lineages as labeled by CLADES 2.0 + GH146-GAL4. (E) Below, number of transitions underwent by different neuronal types (n=63 clones, 49 brains) labeled with GH146. Points with the same color denote the same value. Horizontal lines represent mean. Above, probability (for each neuronal type) of undergoing at least one transition. Points were placed between two neurons to represent the transition occurred between two types. Note the increase during quiescence and in the latest type (VAL1lm). (F) Representative example of a lineage labeled with CLADES 2.0 + Acj6-GAL4 (n=6 clones, 6 brains). Green, red, blue and gray, immunohistochemistry for V5 (YFP), RFP, HA (CFP) and Nc82 respectively. Scale bars = 50 micrometers in B and 15 micrometers in D, F.

Neuroblasts in the larval brain were predominantly labeled with green, yellow and/or red fluorescence and a lower proportion of neuroblasts also reached the purple (RFP+CFP) and blue steps in the cascade. Despite every further step in the cascade is less likely to occur due to the occurrence of incorrect repair outcomes, the proportion of red neuroblasts was higher compared to yellow neuroblasts (Fig. S9B). This is probably due to the fact that, besides via SSA, the reporter inactivation can also occur as a result of incorrect repair events bringing the reporter out of frame (roughly 66% of indels assuming an equiprobable distribution).

To demonstrate the power of CLADES 2.0 for lineage tracing, we set out to reconstruct one of the most heterogeneous lineages in *Drosophila*, the ALad1 lineage. This lineage generates 40 morphologically distinguishable neuronal types in a known developmental sequence (Fig. 4C, 8). To this end, we employed two GAL4 lines, Acj6-GAL4 to label the entire lineage and GH146-GAL4 to selectively mark most of the early-born neuronal types (Fig. 4C). Each GAL4 line was crossed to a fly bearing Dpn-Cas9 and the CLADES 2.0 constructs. Overall, patterns for both GH146 (Fig. 4D-E, Table S3) and Acj6 (Fig. 4F) were consistent with previous lineage studies based on twin-spot MARCM (8). For GH146, we calculated the average number (N=63 ALad1 clones) of cascade steps underwent by the progenitor cell and inherited by each neuronal type (Fig. 4E). Although the exact birth-order of some of the neuronal types (especially in the early window) could not be determined based on this parameter, the progressive increase along the temporal axis allowed us to resolve most part of this lineage. At the single-clone level, the general progression of the cascade also correlates with the previously described birth-order. We only found occasional inconsistencies at the single-cell level in 29% of clones (Table S3). This phenomenon is likely the consequence of Cas9 protein perdurance, and thus cascade progression in the GMC. Interestingly, most of the clones revealed the embryonic and larval-generated neurons in different colors, suggesting an increased probability of cascade progression during the period of neuroblast quiescence (Fig. 4E). In some cases, the cascade even progressed two steps during quiescence (Table S3). In summary, these results demonstrate that CLADES 2.0 allows to reconstruct complex lineages from a limited number of samples.

### CLADES as an event tracker: cascade progression across fly generations

The utility of CLADES is not limited to neuronal lineage progression. If Cas9 activity is expressed under different conditions, CLADES should function as a reporter of other biological events. To explore this application, we set out to use CLADES as a reporter of fly generations (Fig. 5A-E).

**Fig 5.**
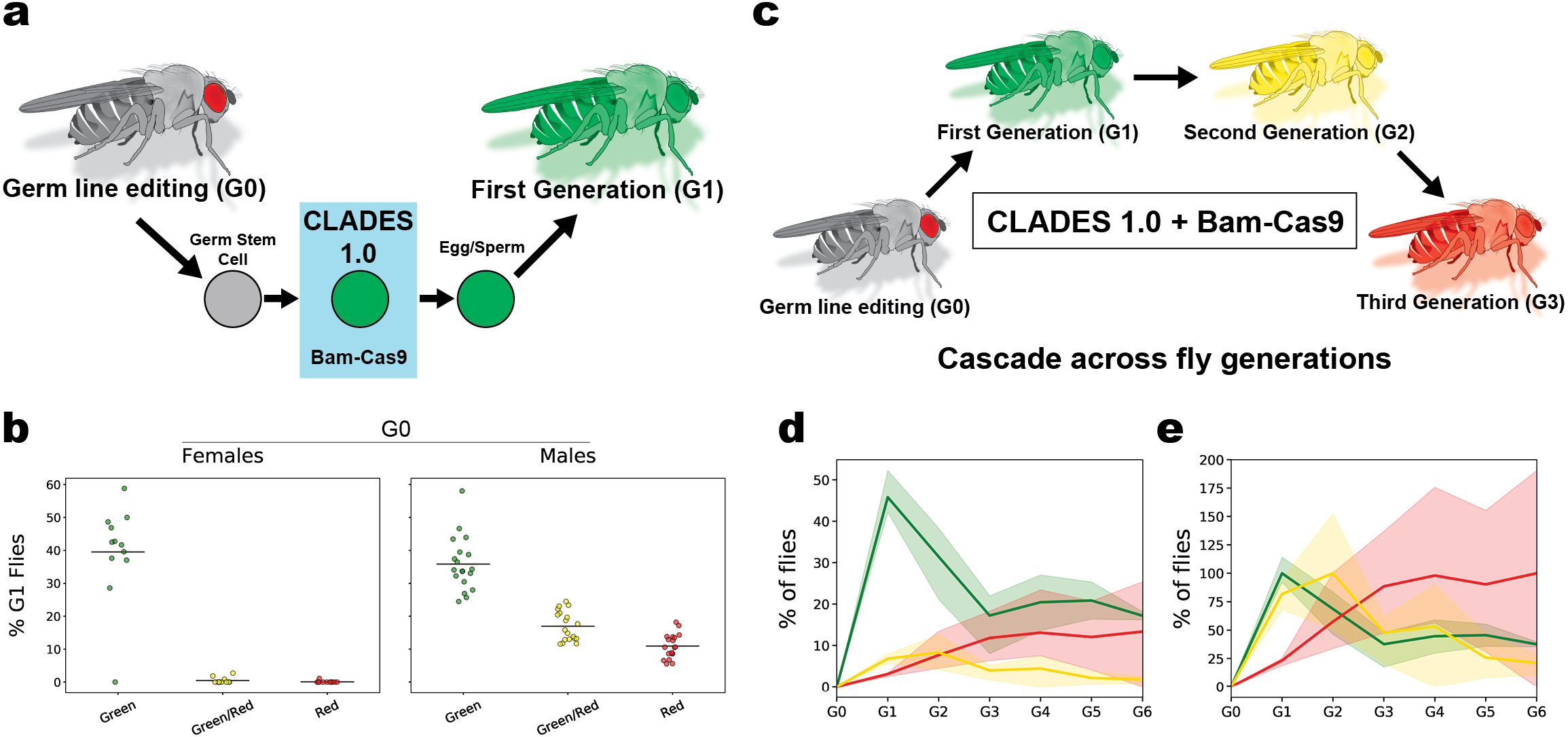
CLADES as an event tracker: cascade progression across fly generations. (A) Cartoon illustrating the progression of CLADES 1.0 in the germ line for the single-generation experiment. (B) Percentage of G1 flies labeled with each reporter combination (n=12 and 18 independent crosses for females and males respectively). Horizontal lines represent mean. (C) Scheme illustrating the progression of CLADES 1.0 across multiple fly generations. (D) Percentage of flies expressing each combination of reporters in each generation (n=3 independent crosses, 20 parents per cross). (E) Percentage of flies normalized to the maximum average value. Areas surrounding the line plot represent the 95% confidence interval.

In order to track multiple generations with limited reporters, restricting the cascade progression once per generation is critical. To this end, we expressed Cas9 with the *bam* promoter, which is transiently active in the germ line when germline stem cell (GSC) progeny initiate differentiation (15–16). Therefore, to test CLADES as a reporter of fly generations, we crossed a fly line expressing Bam-Cas9 and the trigger U6-gRNA#1 to CLADES 1.0. First, we quantified the number of steps the cascade progressed over a single generation (Fig. 5A-B). To accomplish this, we generated flies (G0) expressing the two CLADES reporters, the trigger U6-gRNA#1, and Bam-Cas9, thus initiating the cascade in the germ line. We then crossed individual G0 males and females with double balancer flies and quantified the percentage of G1 progeny expressing each reporter. While progeny from G0 females had consistently advanced a single cascade step, the cascade could progress multiple steps in the progeny of the G0 males (Fig. 5B). Next, we wanted to follow the cascade progression across multiple generations at the population level. For that, we crossed G0 males to G0 females and selected G1 flies bearing only one CLADES allele. We repeated this scheme until the sixth generation (G6). We observed that the cascade progressed as expected (Fig. 5D). While the efficacy decayed after the first step, the final population also contained a higher percentage of red flies compared to yellow flies (as occurred when CLADES 2.0 was targeted to neuroblasts, Fig. S9). We took this decay into account and plotted the normalized cascade progression in Figure 5E. G1 was easily distinguished by the high proportion of green flies and yellow flies, which arises as a result of the combination of gene editing from female and male parents observed in Fig 5B. In G2, the proportion of yellow flies reached its peak (Fig. 5D&E), concurrent with a decrease of green and the increase of red flies. In G3-G4 the proportion of red flies reached its maximum as yellow flies kept decreasing and green flies reached a plateau. Finally, the populations remained seemingly stable in G5-G6. In summary, when applied to the germ line, CLADES makes it possible to track generations of flies (at the population level) based on the proportion of reporters expressed. Also, it allows to delay the activation of reporters/gRNAs for several generations (see discussion).

## Discussion

Synthetic biology holds immense potential for programming complex biology, yet its development has been largely unavailable to multicellular organisms (17). CLADES overcomes this system barrier in *Drosophila* and allows the execution of a sequence of conditional clauses: a next step is triggered only upon activation or deactivation of the previous step. Here we have used this concept for the sequential transgene activation/deactivation. We have thus created three-to five-color reporter cascades, therefore maximizing the number of colors that can be generated from a given number of reporters. This allowed us to reconstruct lineages from the specific reporter combination expressed by each of the daughter cells. Despite the complex system of genetic relays, CLADES 2.0 lineage tracing is simple as it only requires crossing a CLADES line to a GAL4 line of interest.

As a proof-of-concept, we reconstructed the birth order of the ALad1 lineage. We reached a good balance between the resolution and the number of brains required in the analysis. While we could potentially improve the resolution by increasing the number of brains, the resolution is more likely limited by how fast the cascade can progress. The relatively slow progression makes it unlikely to deduce the order of embryonic-born neurons that arise rapidly during the initial phase of fly neurogenesis. Another limitation of CLADES is the perdurance of Cas9 activity in the GMC. Some of the brains in the analysis contained a few neurons whose order was different from previously described (8). In some cases, these GMC events were particularly abundant over certain time windows, leading to multiple neurons innervating the same glomerulus to be labeled in a non-chronological manner. Thus, an accurate reconstruction requires sampling multiple brains. However, the number of brains necessary for accurate lineage reconstruction is extraordinarily low (∼50) when compared to previous methodology requiring thousands of brains (8). Moreover, this issue could be solved by tightly controlling the concentration of Cas9 in the GMC with destabilizing protein domains such as geminin (18). Other issues center around the progressivity of the cascade which can prematurely cease due to unwanted indels. This sub-optimal progressivity makes it unrealistic to substantially increase the number of steps in the cascade or to create multidimensional cascades (cascades that could be triggered by other cascades). However, SSA effectiveness can be as high as 95% when longer repeats are used (9). Therefore, creating gRNAs scaffolds with longer repeats may improve CLADES drastically. Including more than one target site in the switch (so that there is more opportunities for SSA) or increasing the number of copies for each construct could also help to make the expected repair outcome more likely. Future CLADES versions will incorporate these improvements, as well as an increased editing rate (e.g. by increasing Cas9 concentration) to speed the cascade progression for more fine-tuned lineage reconstruction.

In its current form, CLADES creates numerous opportunities. First, it allows to trace entire lineages from a reduced number of brains. Unlike any existent technology, one can also have a lineage divided into five temporal genetic windows in a single animal. This five-colors resolution makes it possible for rapid screening of molecular factors involved in temporal specification, as mutant phenotypes can be associated with birth-order. Further, since Cas9 works efficiently in most species (19), this tool should be readily applicable to other animal models where sophisticated lineage tools are not yet available. In vertebrates, neurons usually undergo migration (20), making morphological identification more straightforward as they do not cluster together. Cell cycles are also generally slower (21), which should reduce the number of neurons labeled with the same reporter(s). Another possibility is creating a cascade of transcriptional activators such as GAL4, LexAp65 or QF (in *Drosophila*). One could thus inhibit (i.e. UAS-RNAi) or overexpress genes (i.e. to induce apoptosis) and report each manipulation with different reporters. CLADES could also be used to differentiate stem cells into specific cell types *in vivo* by delivering genetic cascades. This approach mimics the natural cell specification process that often occurs as cascades of transcription factors (22–24). CLADES can also be combined with inducible forms of Cas9 or gRNAs. In that way, one could report cellular events occurring at a specific time. When applied to the germ line, CLADES may help to optimize the Cas9-based gene drive technology. One of the main limiting factors of this technology is the accumulation of mutations in the gRNA target sequence, thus preventing the spread of the gene drive (25). CLADES could trigger different gene drives in different generations, thus reducing the emergence of resistant alleles. It could also delay the activation of genes that, in turn, could stop the spread of the gene drive after several generations. To our knowledge, CLADES is the only existing technology enabling this type of transgenerational genetic manipulations.

Considering lineage tracing, one future prospect of the CLADES concept excels among all others. The advent of methods based on the progressive accumulation of DNA mutations should, in theory, allow scientists to reconstruct whole-animal lineages (5–7). Most of these methods, such as Gestalt (7), use Cas9 to induce mutations in the DNA. However, the target sites for these mutations become rapidly depleted and only minor portions of lineages can be reconstituted. Combining lineage tracing via accumulated DNA mutations and CLADES could resolve this problem. Cascades of different gRNAs could be triggered with each gRNA introducing mutations in a dedicated region. Thus, those targets in each region would remain unedited until the corresponding gRNA becomes active. Computer simulations showed these gRNA cascades enable the extended progression of mutations and therefore the length of the lineage that can be resolved (26). Moreover, CLADES would allow to follow lineage progression based on imaging. Such tool combination may become the ultimate lineage tracing method to track whole-animal development.

## Acknowledgments

We thank all members of Tzumin’s lab for their comments and feedback, especially Rosa Miyares for critical reading and input on the manuscript. We also thank Haluk Lacin and Eduardo Martin-Lopez for their input on the manuscript. We thank Qingzhong Ren and Janelia Fly Core for their excellent technical support. We thank our suppliers Rainbow, Genscript and Benchling for their services. We thank *Drosophila* Genomics Resource Center, supported by NIH grant 2P40OD010949, for the S2 cell line. Stocks obtained from the Bloomington *Drosophila* Stock Center (NIH P40OD018537) were used in this study. We thank Crystal Di Pietro and Kathryn Miller for administrative support

## Author contributions

Conceptualization, J.G.-M. and T.L.; Methodology, J.G.-M. and T.L.; Investigation, J.G.-M., C.-P.Y., I.E.-M. and K.M.; Writing – Original Draft, J.G.-M and T.L.; Writing – Review & Editing, J.G.-M., C.-P.Y., I.E.-M., M.K. and T.L.; Visualization, J.G.-M., Supervision, M.K. and T.L.; Project Administration, M.L. and T.L.; Funding Acquisition, M.K. and T.L.

## Materials and Methods

### Plasmids Construction

All the DNA constructs were designed with Benchling (Benchling platform) and generated by standard cloning techniques, including restriction digest/ligation and PCR assembly. The sequences for the gRNAs were selected based on their low off-target (27) and their high on-target activity (28). The lowest score was 79.8 and 93.9 for on-target and off-target activity respectively (see Table 2). The final constructs were verified by sequencing.

#### Conditional gRNA

- Variant #1: a FseI-HH-gRN(#2)NA#1-HDV-HindIII fragment was *de novo* synthetized (<u>Genscript)</u>. This contains a conditional gRNA#1 activatable by the gRNA#2 and flanked by the Hammerhead and HDV ribozymes (29). This fragment was then cloned into a FseI/HindIII site in DpnEE-pBPKD1Uw (Tzumin Lee lab).
- Variant #2: a block containing the conditional U6:3 promoter (activatable by the gRNA#2) and the gRNA#1 was *de novo* synthetized (gBlock, IDT) and cloned into a HindIII-EcoR site in pCFD3 (30)
- Variants #3-6: a cassette containing the U6:3 promoter (30) and the corresponding gRN[#1]NA#2 was assembled by PCR and then cloned into a HindIII-EcoRI site in pCFD3.
- Variants #7-12: a block containing the U6:3 promoter and the corresponding gRN(#2)NA#1 was *de novo* synthetized (gBlock, IDT) and cloned into a HindIII-EcoR site in pCFD3.
- Variants #13-14: a cassette containing the U6:3 promoter and the conditional gRN(#2)NA#1 was assembled by PCR. This step included PCR-amplifying part of the Neomycin gene as an arbitrary sequence to increase the length of the repeats involved in SSA. The full cassette was then cloned into a HindIII-EcoR site in pCFD3.

#### CLADES

- Construct #1: we first designed this construct so that the reporter is initially out of frame. After the ON cassette is activated, the reporter becomes on frame and none stop codon should exist in the sequence. To that end we selected the ORF with fewer stop codons and added random nucleotides when necessary. We also tailored the U6 promoters minimizing the number of modifications, especially in the regulatory regions (PSEA and TATA box, 31). To remove stop codons and decrease the occurrence of repeats within the construct, we also modified the gRNA scaffolds without altering the secondary structure (32). We then generated an EagI-myr::mCherry-p10-EagI fragment by PCR amplifying from 10XUAS-IVS-myr::mcher[#1]rry (9). Given the presence of large repeats in the mcher[#1]rry gene, the PCR produces the reconstituted mCherry as a subproduct. This fragment was then cloned into Actin5C-IVS-myr::GF[#2]FP (9), thus generating Actin5C-IVS-mCherry. We next assembled a fragment NheI-U6:2-gRN(#5)NA#2-T2A::myr-BamHI (OFF cassette+T2A::myristoylation signal) by PCR. This contained a U6:2 promoter (31) driving a conditional gRNA#2 (activatable by gRNA#5), the sequence for the T2A peptide (to release the reporter from the OFF cassette) and a myristoylation signal (to direct the reporter to the plasmatic membrane). We also assembled a XhoI-U6:3-gRN(#3)NA#1-NheI fragment (ON cassette) by PCR. Both fragments were then cloned into the XhoI/BamHI site in Actin5C-IVS-mCherry. The conditional gRNAs used for this plasmid are not described in Figure S1. Hereon, all the CLADES plasmids contain the variant #6 for all the conditional gRNAs.
- Construct #2: we generated a mCherry version in which the first four ATG codons were substituted, still exhibiting strong fluorescence. Briefly, we used degenerate primers to PCR-amplify mCherry from Actin5C-IVS-mCherry. These primers contained the degenerate sequence NNH instead of the ATG codons. The PCR product was then cloned into a bacterial vector for protein expression (pJet1.2/blunt; Thermo Fisher Scientific) and transformed in bacteria. Subsequently, we sequenced the colonies with the strongest fluorescence. We finally selected the variant M1L, M10P, M17Q and M23C that we named mPicota (after a stemless cherry variety uniquely grown in Cáceres, Spain). Next, a XhoI-U6:3-gRN(#3)NA#4-NheI (ON cassette) and a NheI-U6:2-gRN(#5)NA#6-T2A::myr-BamHI cassette (OFF cassette+T2A::myristoylation signal) were assembled by PCR. In these cassettes, two ATG codons in the myristoylation signal and one in the U6:2 promoter were removed to avoid translation starting after the ON cassette. These fragments and a BamHI-mPicota-p10(1-44bp)-BsiWI were then cloned into a XhoI-BsiWI site in the Construct #1.
- Construct #3: we sought to remove all the predicted splicing donor and acceptor sites from the Kozak sequence to the beginning of the reporter. We thus designed a sequence where the highest score was 0.26 (NNSPLICE 0.9, 33). We introduced multiple modifications along the sequence, following the guidelines described for the construct #1. To introduce these modifications, we assembled a cassette KpnI-IVS-U6:3-gRN(#1)NA#3-U6:2-gRN(#4)NA#7-T2A-myr-BamHI by PCR. With a future cascade in mind, we changed the reporter and generated a BamHI-3XV5-mCitrine-BsiWI cassette by PCR. Both were then cloned into a BamHI/BsiWI site in the Construct #1.
- Construct #4: a XhoI-U6:3-gRNA#3-SapI (preactivated ON cassette) cassette was cloned into a XhoI/SapI site in the Construct #3.
- Construct #5: a XhoI-SapI fragment containing the Kozak sequence and first start codon was assembled by primer annealing. We then cloned it into a XhoI/SapI site in the Construct #4, thus removing the ON cassette.
- Construct #6: a SapI-T2A::myr::3XV5-BamHI fragment was assembled by PCR and cloned into a SapI/BamHI site in the Construct #4, thus removing the OFF cassette. To reduce the plasmid size and thus increase the probability of transfection, we removed the mini-white marker by AscI digestion and re-ligation.
- Construct #7: we generated (by PCR) a KpnI-IVS-U6:3-gRNA#3-SapI cassette in which the Kozak sequence and start codon were moved to the end of the U6 promoter and two ATG codons on frame located in the U6:3 promoter were removed. This was cloned into a KpnI/SapI site in the Construct #6 (before removing the mini-white marker).
- Construct #8: a SapI-T2A-U6:2-gRN(#4)NA#7-T2A-myr-3XV5-BamHI cassette was generated by PCR and cloned into a SapI/BamHI site in the Construct #7. Finally, we removed the mini-white marker by AscI digestion and re-ligation.
- Construct #9: we generated a SapI-spacer-U6:2-gRN(#4)NA#7-T2A-myr-3XV5-BamHI cassette by PCR and cloned it into a SapI/BamHI site in the Construct #8.
- Construct #10: we first designed a U6:3 promoter without the PSEA and TATA-box sequences. This was assembled into a XhoI-U6:3-gRNA#3-SapI cassette by PCR and cloned into a XhoI/SapI site in the Construct #6.
- Construct #11: we designed a construct with controlled splicing in which a splicing donor (score>0.90) was introduced immediately upstream of each U6 promoter. Similarly, a splicing acceptor was embedded in the final region of each U6 promoter. This construct also contains the Kozak sequence and the starting codon at the end of the U6:3 promoter. To clone it, we assembled two cassettes by PCR, a KpnI-U6:3-gRNA#3-SapI and a SapI-U6:2-gRN(#4)NA#7-T2A-myr-3XV5-BamHI. These fragments were then inserted into a KpnI/BamHI site in the Construct #3.
- CLADES1.0-YFP (Construct #12): the final CLADES design was generated by refining the Construct #11. First, we restored the original sequence of the U6 promoters in those modifications aimed to remove stop codons. Due to the controlled splicing, those modifications were no longer necessary as those regions will not be part of the mRNA. These modifications generated a new ATG codon on frame in the U6:2 promoter that we removed by modifying a single nucleotide. Similarly, we moved back the Kozak sequence and the start codon upstream of the ON cassette. We also introduced a strong splicing acceptor before the Kozaq sequence, aimed to capture any potential endogenous splicing occurring upstream. Finally, we removed the myristoylation signal and introduced a CAAX signal at the end of the reporter. This was codon optimized for *Drosophila*, as well as the T2A, V5 and CAAX sequences. Following this design, we synthetized three fragments (gBlock, IDT): i) a KpnI-U6:3-gRN(#1)NA#3-SapI, ii) a SapI-U6:2-gRN(#4)NA#7-NotI and iii) a NotI-T2A::3XV5::mCitrine::CAAX-BsiWI. These were cloned into a KpnI/BsiWI site in the Construct #1.
- CLADES1.0-RFP: following the same design used for the Construct #12, three fragments were synthetized (gBlock, IDT): i) a KpnI-U6:3-gRN(#3)NA#4-SapI, ii) a SapI-U6:2-gRN(#5)NA#6-NotI and iii) a NotI-T2A::mPicota::CAAX-BsiWI. These were cloned into a KpnI/BsiWI site in the Construct #1.
- CLADES1.0-CFP: as explained for Construct #12, we synthetized three fragments (gBlock, IDT): i) a KpnI-U6:3-gRN(#7)NA#5-SapI, ii) a SapI-U6:2-gRN(#8)NA#9-NotI and iii) a NotI-T2A::3XHA-mTurquoise2::CAAX-BsiWI. These were cloned into a KpnI/BsiWI site in the Construct #1.
- CLADES2.0 (CFP, YFP, RFP): an EcoRI-10XUAS-hsp-KpnI cassette was cloned into an EcoRI/KpnI site in the corresponding CLADES1.0 construct.

#### Triggers

U6-gRNA#3 and U6-gRNA#4: annealed primers with the corresponding spacer (the part in the gRNA providing the specificity) were cloned into a SapI site in pCFD3.

#### Zebrafish

- Ubi-SpCas9::P2A::mPicota::T2A::mCitr(#1)trine-polyA-U6c-gRN(#7)NA#1: we used multisite-Gateway cloning to recombine: i) a p5E *ubi* vector (34, Addgene #27320), ii) a pME vector carrying SpCas9-P2A-mPicota-T2A-mCitr(#1)trine-polyA (9), iii) a p3E vector containing U6c-gRN(#7)NA#1 and iv) a pDestTol2 vector (35). The p3E vector was built by cloning a U6c-gRN(#7)NA#1 fragment (gBlock, IDT) into a KpnI/XhoI site in p3E-MCS (Addgene #75174).
- Ubi-Cas9-P2A-mPicota-T2A-mCitr(#1)trine-polyA-U6c-gRN(#7)NA#1-U6d-gRNA#7: we first generated p3E-U6c-gRN(#7)NA#1-U6d-gRNA#7 by cloning a fragment U6c-gRN(#7)NA#1 into a KpnI/XhoI in p3E-U6c-gRNA#7 (p3E-U6c-gRNA#3 in 9). Then, the p3E vector was used for the gateway reaction explained above.

### Generation of transgenic flies

Most lines were generated by using the PhiC31 system (36). Injections were performed by Rainbow Transgenic Flies Inc.

Those CLADES lines with a preactivated ON or OFF cassette were generated by crossing the original CLADES line with a fly bearing Actin-Cas9 and the corresponding gRNA under U6 promoter. Stocks from 10 G1 flies were established and screened by Sanger sequencing to confirm a perfect SSA event.

### Fly strains

#### Previous stocks

The following lines were used in this study: Actin5C-Cas9 (ZH-2A, BDSC#54590), w; Sp/CyO; MKRS/TM6B (Janelia Fly Facility), BamP-Cas9 (Attp2, Tzumin Lee lab), GH146-GAL4 (BDSC#30026), Acj6-GAL4 (BDSC#30025) and DpnEE-GAL4 (Attp16, 37). The rest of lines were generated in Garcia-Marques et al. (9): GR44F03-3XgRNA#1 (Attp40), Actin5C-IVS-myr::GF[#2]FP (VK00018), Actin5C-IVS-myr::mcher[#1]ry (Attp2), DpnEE-IVS-Cas9 (VK00033), DpnEE-IVS-Cas9 (Attp40), DpnEE-IVS-Cas9 (Attp2), U6-gRNA#1 (Attp2) and U6-gRNA#1&2 (Attp40).

#### Stocks generated

The following lines were generated for this work: CLADES1.0-RFP(VK00033), U6-gRNA#3 (Attp40), U6-gRNA#4 (Attp40), Dpn-gRN[#1]NA#2 (Variant #1)(Attp40), U6-gRN(#2)NA#1 (Variant #2) (Attp2), U6-gRN[#1]NA#2 (Variant #3)(Attp2), U6-gRN[#1]NA#2 (Variant #4)(Attp40), U6-gRN[#1]NA#2 (Variant #5)(Attp2), U6-gRN[#1]NA#2 (Variant #6)(Attp2), U6-gRN(#2)NA#1 (Variant #7) (Attp2), U6-gRN(#2)NA#1 (Variant #8) (Attp2), U6-gRN(#2)NA#1 (Variant #9) (Attp2), U6-gRN(#2)NA#1 (Variant #10) (Attp2), U6-gRN(#2)NA#1 (Variant #11) (Attp2), U6-gRN(#2)NA#1 (Variant #12) (Attp2), U6-gRN(#2)NA#1 (Variant #13) (Attp2), U6-gRN(#2)NA#1 (Variant #14) (Attp2), CLADES_optim_const #1 (Attp40), CLADES_optim_const #2 (Attp40), CLADES_optim_const #3 (Attp40), CLADES_optim_const #4 (Attp40), CLADES_optim_const #7 (Attp40), CLADES_optim_const #8 (Attp40), CLADES_optim_const #11 (Attp40), CLADES_optim_const #12 (Attp40, also referred to as CLADES1.0-YFP), CLADES1.0-YFP(OFF pre-activated) (Attp40), CLADES1.0-YFP (VK00020), CLADES1.0-RFP(Attp2), CLADES1.0-RFP(VK00033), CLADES1.0-RFP(ON pre-activated) (VK00033), CLADES1.0-RFP(ON and OFF pre-activated) (VK00033) and CLADES1.0-CFP (VK00027).

### S2 culture and transfections

*Drosophila* S2 cells (S2-DGRC) were grown in Schneider’s *Drosophila* Medium (Thermo Fisher Scientific) containing 10% heat inactivated fetal bovine serum (Thermo Fisher Scientific) and pen/streptomycin (Thermo Fisher Scientific). Transfections we were performed with the Effectene Transfection Reagent (Qiagen), following the manufacturer’s instructions.

### Zebrafish injections

Adults (3 months-2 years old) were mated to generate embryos. Tol2 mRNA was synthetized from linearized plasmid using the mMessage mMachine SP6 Transcription kit (Thermo Fisher Scientific) and purified (RNAeasy Mini Kit, Qiagen) before injection. About 400 embryos for each experiment were injected at 1-cell-state with 1-2 nanoliters of 25 ng/ul of Tol2 transposase mRNA and 25 ng/ul of the corresponding Tol2-conditional gRNA plasmid. Fluorescence was examined after 1 dpf.

### Immunostaining and Antibodies

Larval and adult brains were dissected in PBS and fixed with 4% paraformaldehyde (Electron Microscopy Sciences) in PBS for 40 minutes. Samples were washed three times in PBS containing 0.5% Triton X-100 (Thermo Fisher Scientific) and then incubated at 4°C overnight with a solution of primary antibodies, diluted in PBS containing 5% normal goat serum (Thermo Fisher Scientific). We used the following primary antibodies: anti-Bruchpilot (1:50; DSHB), rat anti-Deadpan (1:100; Abcam), mouse anti-HA (1:500; Roche), rabbit anti-V5 (1:500; Abcam), rat anti-RFP (1:500; Chromotek), rabbit anti-DsRed (1:500; Clontech) and rat anti-GFP (1:500; Nacalai).

After rinsing three times in PBS, samples were incubated overnight at 4°C with secondary antibodies (diluted 1:1000 in PBS). The secondary antibodies used were DyLight 405 AffiniPure Goat Anti-Mouse IgG (Jackson ImmunoResearch), Alexa Fluor® 647-AffiniPure Goat Anti-Mouse IgG (Jackson ImmunoResearch), Alexa 488-conjugated goat anti-Rat IgG (Thermo Fisher Scientific), Alexa 488-conjugated goat anti-Rabbit IgG (Thermo Fisher Scientific), Alexa 568-conjugated goat anti-Rabbit (Thermo Fisher Scientific), Alexa 568 goat anti-Rat IgG (Thermo Fisher Scientific) and Alexa 647 goat anti-Mouse IgG (Thermo Fisher Scientific). Finally, brains were washed again in PBS and mounted using SlowFade^TM^ Gold Antifade Mountant (Thermo Fisher Scientific).

### Image acquisition and processing

Samples were imaged on a Zeiss LSM 880 confocal microscope and processed with Fiji(NIH) and Adobe Photoshop CC 2018 (Adobe).

For zebrafish imaging, animals were anesthetized by bath application of 0.02% w/v solution of Ethyl-3-aminobenzoate methanesulfonate (Sigma-Aldrich, St. Louis) in filtered fish system water for 1 min. Fish were then mounted in a drop of 1.6% low melting point agarose (Invitrogen) over a glass-bottomed plate.

For imaging whole-flies, flies were frozen and adhered to slides with vacuum grease(EMS, cat#60705). We then imaged slides with a stereo fluorescence microscopy (Olympus). We did not quantify CFP+ flies given the technical impossibility of distinguishing real signal from autofluorescence in whole CFP+ flies.

### Analysis of repair outcome

Flies were anesthetized with CO_2_ and the head was dissected with forceps. Genomic DNA from 30 heads (for each replicate) was extracted by using the DNeasy Blood & Tissue kit (Qiagen), following the manufacturer’s instructions. DNA samples (200 ng) were used in PCR reactions to amplify the region of interest, using the Q5 2X Master Mix kit (New England Biolabs). PCR reactions were performed as indicated by the manufacturer, with 72 degrees as the melting temperature. The primers (with partial Illumina adapters) used were:

Amplicon_F:ACACTCTTTCCCTACACGACGCTCTTCCGATCTNNNNNNCGCCAAGCAGAGAGGGCGCCAGTGCTC

Amplicon_R: GGACTGGAGTTCAGACGTGTGCTCTTCCGATCTCAAAAAAAGCACCGACTCGGTGCC AC

Amplicons (∼300 bp) were gel-purified using the QIAquick Gel Extraction kit (Qiagen). Samples were then sent for NGS-sequencing (Amplicon EZ, Genewitz), recovering a minimum of 50000 reads per sample. Sequencing data was analyzed with a custom algorithm (described below and available on GitHub).

### Quantification and Statistical Analysis

#### Neuroblast counting

A minimum of 10 larval brains were dissected and immunostained to detect Dpn, V5(YFP), RFP and HA (CFP, Figure 8S). We randomly sampled a minimum of 30 dpn+ cells (neuroblasts) from the central brain and ventral nerve cord. We counted the number of neuroblasts expressing each combination of marker and expressed it as a percentage out of the total number of sampled neuroblasts.

#### Algorithm for the analysis of SSA

Amplicon sequencing reads were first checked to remove ultra-short (<30bp) reads as well as reads with undetermined bases (N) or very poor (<4std) averaged sequencing scores. Paired reads were then merged through matching the terminal one fifth of the read1 or read2 sequence with the entire read2 or read1 sequence. Only successfully merged reads were subjected to the following analysis. First, merged reads were clustered based on complete nucleotide sequence to reveal discrete amplicon sequences and their read counts. Second, because the coexisting trigger and target transgenes can be equally amplified, we further retrieved those target-derived amplicon reads based on the target-specific spacer sequence. Third, we aligned each target-derived unique sequence to the target’s amplicon reference to recover a minimal number of non-overlapping perfectly matched segments (>4bp in length) that jointly cover as many common bases as possible. Gap-free sequences were deemed as wild-type reads. Fourth, we searched for gaps possibly resulting from fusion of direct repeats and consistently annotated such gaps with all the originally repeated bases lying on the 3’ side. Fifth, we clustered indels based on the indices of the involved bases. Finally, we chose those indels uncovering any of the 5 bases around the trigger-dependent Cas9 cut (2-6bp away from PAM) for quantification of trigger-induced SSA or other indel events across different samples.

#### Glomerular annotation

Glomeruli were annotated as previously described (38). Clones with only green cells were discarded for the analysis. The average number of steps in the cascade was calculated based on the color of each glomerulus in each ALad1 clone (green=0 steps, yellow=1, red=2, purple=3, blue=4). The transition probability was calculated for each neuronal type by dividing the number of clones in which the cascade progressed in that glomerulus (with respect to the previous glomerulus) by the total number of ALad1 clones (N=63 clones). As Acj6-GAL4 is also expressed in the olfactory receptor neurons, for Acj6-GAL4 glomerular annotation the antennae and the maxillary palps of newly eclosed adult flies were surgically removed. This allows the axons of olfactory receptor neurons to degenerate so that the morphologies of antennal lobe neurons can be distinguished.

## Supplementary Text

### Scaffold optimization (see Fig. S1)

**Variant #1:** the first variant tested was designed to be expressed under type II promoter (most tissue-specific promoters). This required the incorporation of ribozymes to process the gRNA out of the mRNA (29). Assuming this requirement, we hypothesized that each part of the conditional gRNA (before and after the target sequence) would lack the upstream or downstream ribozyme, which would abolish its activity. However, we observed leaky activity even with the more restricted Dpn-Cas9. We also abandoned the idea of implementing conditional gRNAs for type II promoters as these are weaker compared to U6 promoters (data not shown).

**Variant #2**: instead of inactivating the gRNA, this strategy was based on the inactivation of the U6 promoter by introducing a switch between the promoter regulatory elements: PSEA and TATA box. The distance between these elements is fixed and minor modifications have a strong effect on the promoter activity (39). After the induction of SSA in the switch, the repair outcome would restore this distance, thus activating the U6 promoter. Unexpectedly, the U6 promoter exhibited a strong leaky activity even in the initial state.

**Variants #3-8**: given the strict requirements of the gRNA structure (32), we hypothesized that incorporating the switch in the scaffold region should alter the gRNA secondary structure and most likely abolish its activity. Only after SSA repair, the native sequence would be restored. We observed considerable leaky activity for most variants, especially with the ubiquitous Actin-Cas9. Note that variant #5 even exhibited activity for the target sequence, which was also used as a spacer. To prevent this potential activity, we reversed the target sequence from Variant #8 onwards. However, even without this modification, the Variant #6 showed only minor leaky activity with Actin-Cas9 and virtually no leaky activity with Dpn-Cas9 (only one clone per 10 brains).

**Variants #9-12**: in previous designs, each part of the gRNA should not be enough for activity: the region upstream from the target lacks most part of scaffold and the region downstream lacks the spacer (which provides the gRNA specificity). Therefore, we hypothesized that the leaky activity could be originated from the interaction between both parts. To reduce this possibility, we designed new variants incorporating a STOP signal (TTTTTT), which should prevent the downstream sequence from being transcribed. This would limit any potential interaction. Yet, we observed leaky activity in all these variants, suggesting that the STOP signal fails to end part of the transcription, and this is enough for the gRNA activity.

**Variants #13-14**: based on the work of other labs (40) and our own results, gRNAs can retain strong activity even if sequence is appended upstream from the spacer. In an attempt to reduce leakiness and concurrently increase the SSA efficiency (by increasing the length of the direct repeats in the switch), we introduced an accessory sequence in the switch, upstream from the spacer region. Surprisingly, we observed strong leaky expression even when the switch upstream from the spacer was more than 400 bp and contained a STOP signal.

### CLADES optimization

**1st design:** the first design of CLADES was based on the incorporation of ON and OFF cassettes upstream from the reporter gene (construct #1). To avoid direct repeats that otherwise could interfere with the SSA expected outcome, we used two different U6 promoters: U6:3 and U6:2. We also diversified each gRNA scaffold based on previous guidelines (32). In the initial configuration, translation should stop at the ON cassette, therefore not reaching the reporter. To our surprise, flies bearing this construct in the initial configuration (before crossing to Cas9/trigger gRNA) exhibited strong fluorescence. We reasoned that translation might start from ATG codons in the reporter, downstream from the OFF cassette. To prevent this hypothetical translation, we created a reporter version lacking the first ATGs (construct #2). However, flies bearing this construct also exhibited strong fluorescence. We then hypothesized that the existence of cryptic splicing sites upstream from the reporter could change the ORF an bring the reporter on frame even before Cas9 acted. Based on *in silico* prediction, we found strong splicing sites in the U6 promoters. After removing these sites (construct #3), we observed no fluorescence. Moreover, crossing these flies with the appropriate conditional reporter showed the conditional scaffolds being part of the ON/OFF cassettes were still functional even after being embedded into the main gene.

**2nd design:** removing all cryptic splicing sites was necessary to avoid reporter expression in the initial state. However, a pre-activated version (in the ON cassette) of the construct #3 failed to express the reporter (construct #4). To understand the reason for the lack of expression, we interrogated this construct by modifying different parts and analyzed fluorescence in S2 cells or flies. First we hypothesized that the presence of the ON or OFF cassette might interfere with the transcription or translation of the main gene. We therefore generated new constructs by removing the ON (construct #5) or the OFF cassette (construct #6), each as a pre-activated version (reporter on frame). In both cases we did not observe any fluorescence. We also designed a construct (construct #7) where the OFF cassette was removed and the starting codon (and Kozak sequence) was moved downstream from the ON cassette, immediately upstream from the reporter gene (on frame). Even in this configuration the reporter failed to be expressed. Additional efforts in this direction sought to improve transcription/translation by inserting a T2A (construct #8) or a random spacer sequence (construct #9) between the ON and OFF cassettes (in the pre-activated version of the construct). In both cases we failed to achieve the reporter expression. Our second hypothesis was that the transcription from the U6 promoters impeded transcription or translation of the reporter gene. To rule out this possibility, we removed the OFF cassette and all the regulatory elements in the U6 promoter of the ON cassette (construct #10). No significant fluorescence was observed in this construct (reporter on frame). Based on previous work, we hypothesized that splicing could be necessary for transcription (41). Therefore, removing all splicing sites could have abolished the reporter translation. To test this hypothesis, we generated a pre-activated version of a construct (construct #11) with controlled splicing. In this construct the splicing sites where placed at the beginning and the end of the U6 promoters. Besides trying to induce splicing, this design sought to exclude the U6 promoters from the mRNA since these were not necessary for reporter expression and might affect translation or mRNA stability. This construct exhibited strong fluorescence and it was the basis for the final optimization.

**3rd design:** we demonstrated a design based on controlled splicing exhibited strong fluorescence. However, the reporter protein was not anchored to the plasma membrane as expected. We reasoned that the myristoylation signal probably required to be located in the N-terminus of the protein (42). Given the presence of the T2A signal, placing the myristoylation signal in the N-terminus would fail to deliver the reporter protein to the plasma membrane. Therefore, we tested CAAX (43) as an alternative for plasma membrane targeting. This tag was very efficient for membranous labeling, completing the optimization of CLADES. The final construct met all the requirements: i) minimal background in the initial state., ii) activation of the ON cassette produced very strong fluorescence delivered to the plasma membrane and iii) fluorescence was removed after the activation of the OFF cassette.

**Fig. S1.**
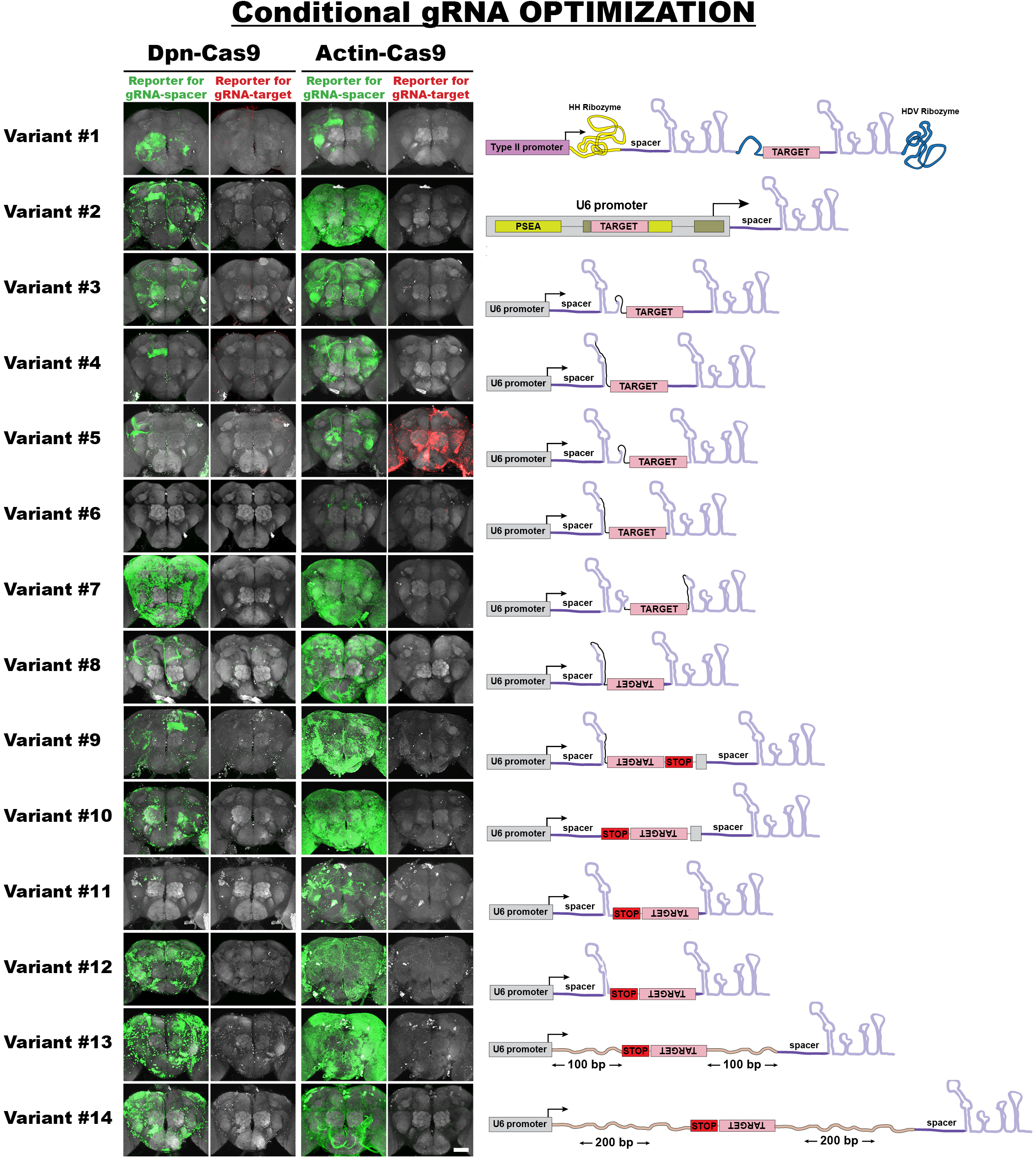
Optimization of a conditional gRNA. 14 conditional gRNA variants were tested for potential leakiness (activity in the absence of the trigger gRNA). We examined the ability of the conditional gRNA to activate, via SSA, a specific conditional reporter (GFP or mCherry) in the absence of the trigger gRNA. Each variant was tested with Dpn-Cas9 or Actin-Cas9. Given that the target sequence in the variant #5 unexpectedly acted as an active gRNA, we included a second reporter (red) to analyze this activity. From the variant #8 onwards we avoided this activity by inverting the orientation of the target sequence. Note that variant #6 is the least leaky as it shows little activity with Actin-Cas9 and almost no activity at all with Dpn-Cas9. Green/Red, immunohistochemistry for EGFP/mCherry. Gray, nc82 counterstain. See also Supplementary Text. N=10 brains. Scale bar = 50 μm.

**Fig. S2.**
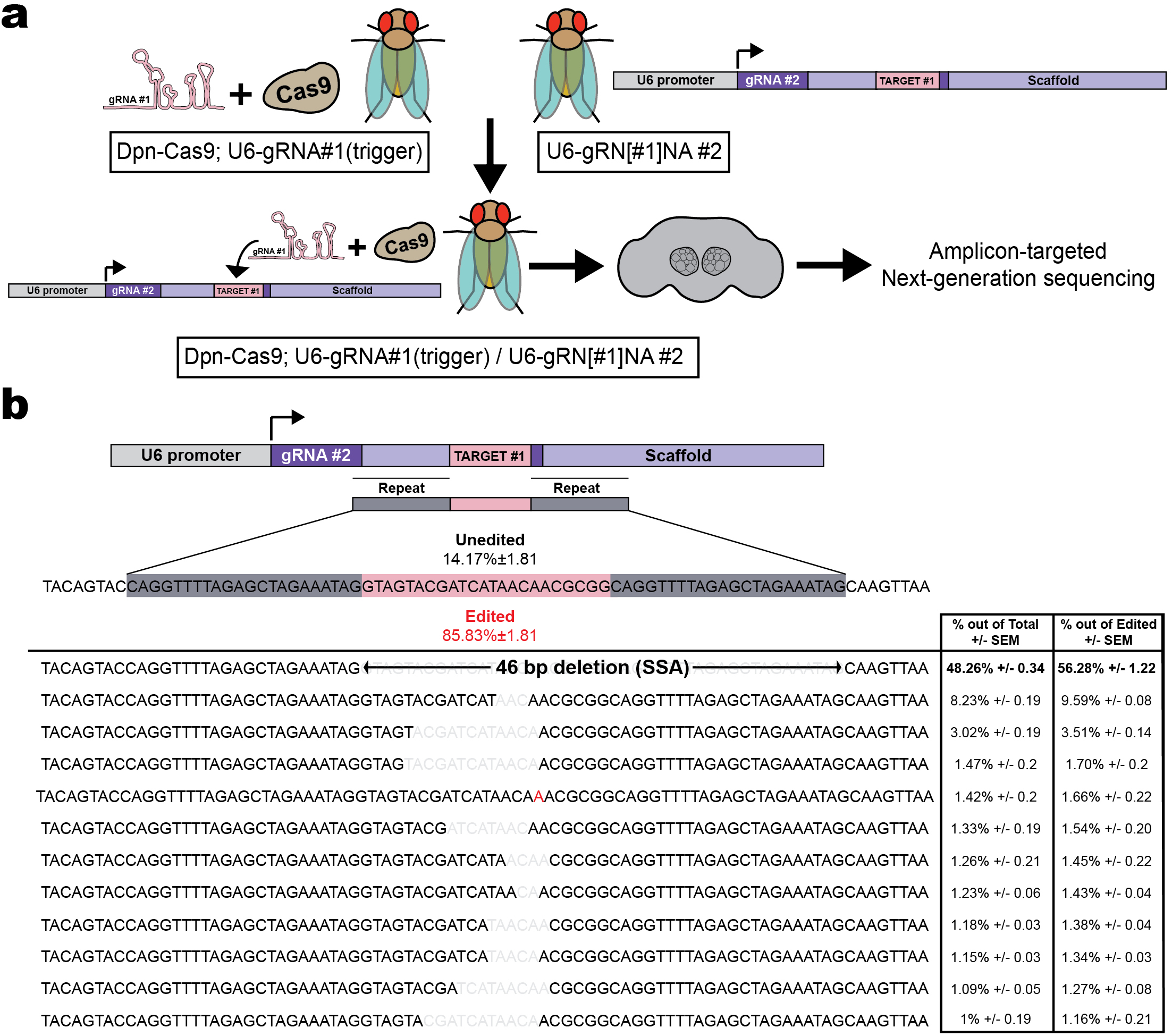
Analysis of the DNA repair outcome by Next-Generation Sequencing. (A) Flies bearing Dpn-Cas9 and the trigger gRNA#1 driven by the U6 promoter were crossed to a fly with the conditional U6-gRN(#1)NA#2. Flies with the three components were analyzed by PCR-amplifying the region flanking the target site and sequencing this amplicon by Amplicon Targeted NGS. (B) Most frequent repair outcomes, including SSA that covers about half of the reads. N=3 replicates, 30 fly heads per replicate.

**Fig. S3.**
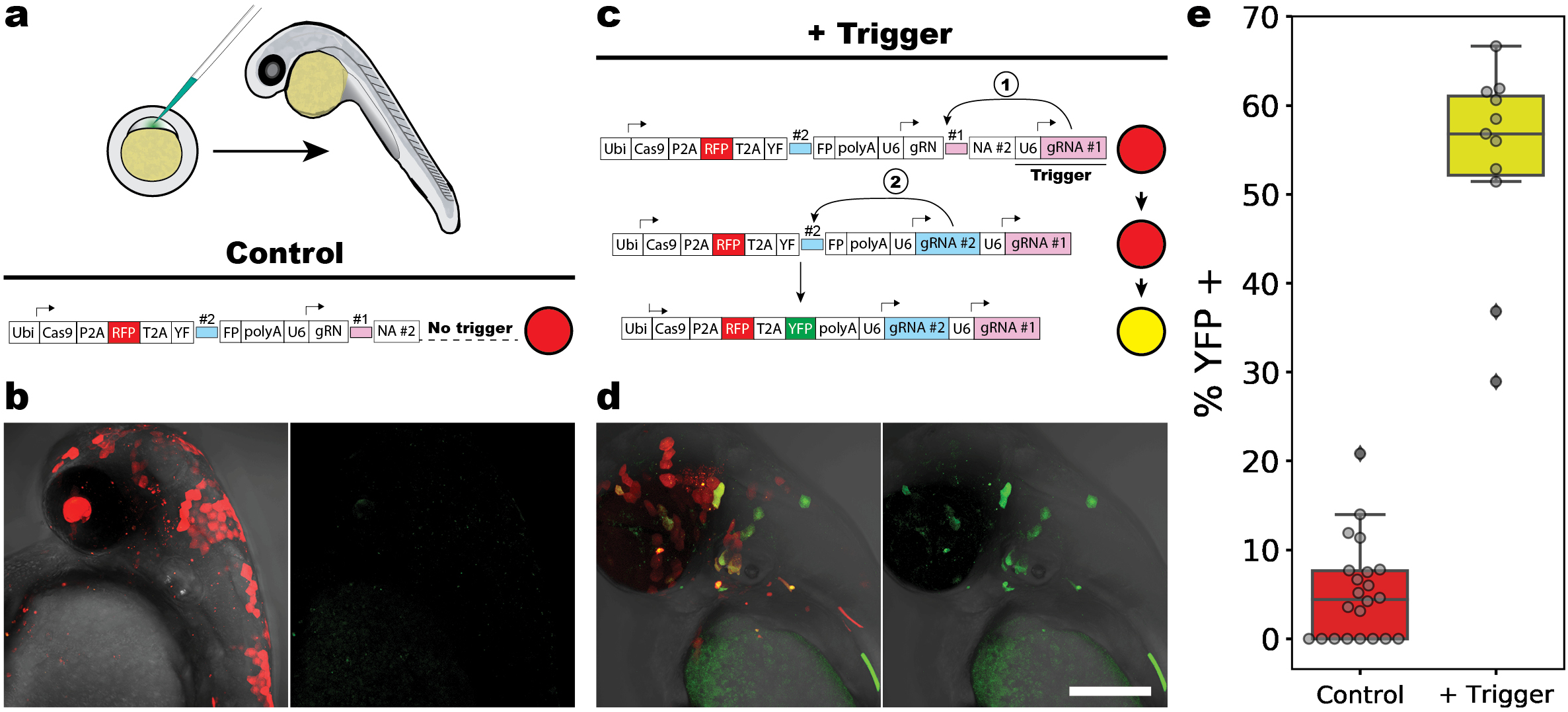
A conditional gRNA works efficiently in a model vertebrate (zebrafish). (A) Tol2-plasmids were injected into 1-cell-stage zebrafish embryos along with mRNA for the Tol2 transposase. All plasmids encoded for: i) Cas9, ii) an RFP protein (injection control) and iii) a YF[#2]FP reporter for the gRNA#2 activity (9). These three proteins shared the same ORF and were under the regulation of the ubiquitous promoter Ubi. Downstream of this cassette we also placed the conditional U6-gRN[#1]NA#2. (B) In the absence of the trigger gRNA, only few cells expressed YFP. (C-D) Adding a U6-gRNA #1 to the control plasmid triggered a gRNA cascade, resulting in most cells expressing YFP. (E) Percentage of YFP+/RFP+ cells. N=22 (control) and 11(experimental), 3 independent experiments. Scale bar = 200 μm.

**Fig. S4.**
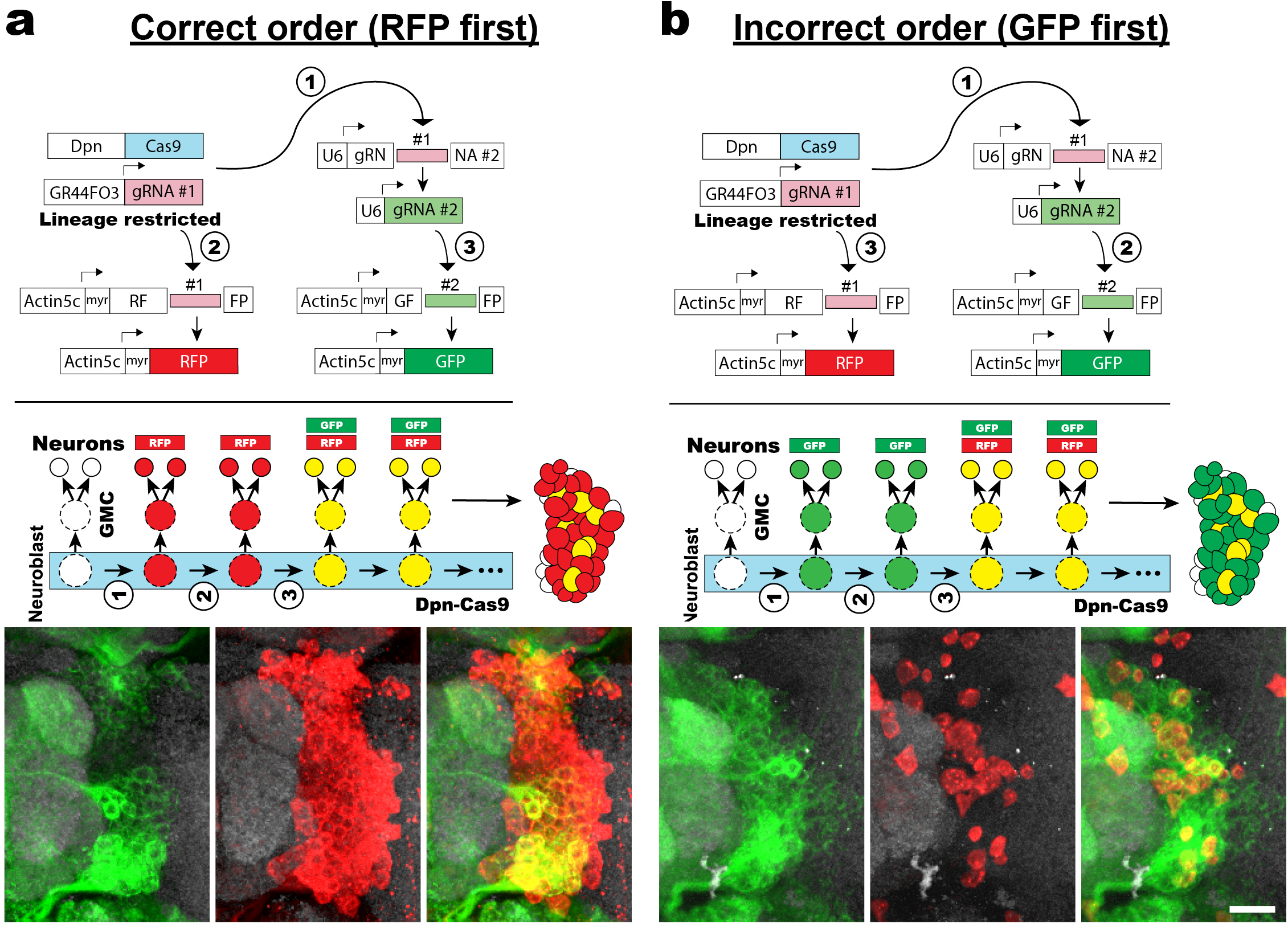
Uncoupling between the gRNA cascade and the reporter cascade. (A) Representative example of the correct order for a cascade of gRNAs controlling the activation of multiple reports *in trans*. Since the gRNA#1 is active from the beginning and the gRNA#2 requires to be activated by the gRNA#1, the activation of the RFP reporter should precede the activation of the GFP. (B) Example where the activation of the second reporter occurs before the first reporter. In both cases the example corresponds to the lateral lineage in the antennal lobe, although this was also observed in other lineages. Green, red and gray, immunohistochemistry for GFP, RFP and nc82 (counterstaining) respectively. Scale bar = 15 μm.

**Fig. S5.**
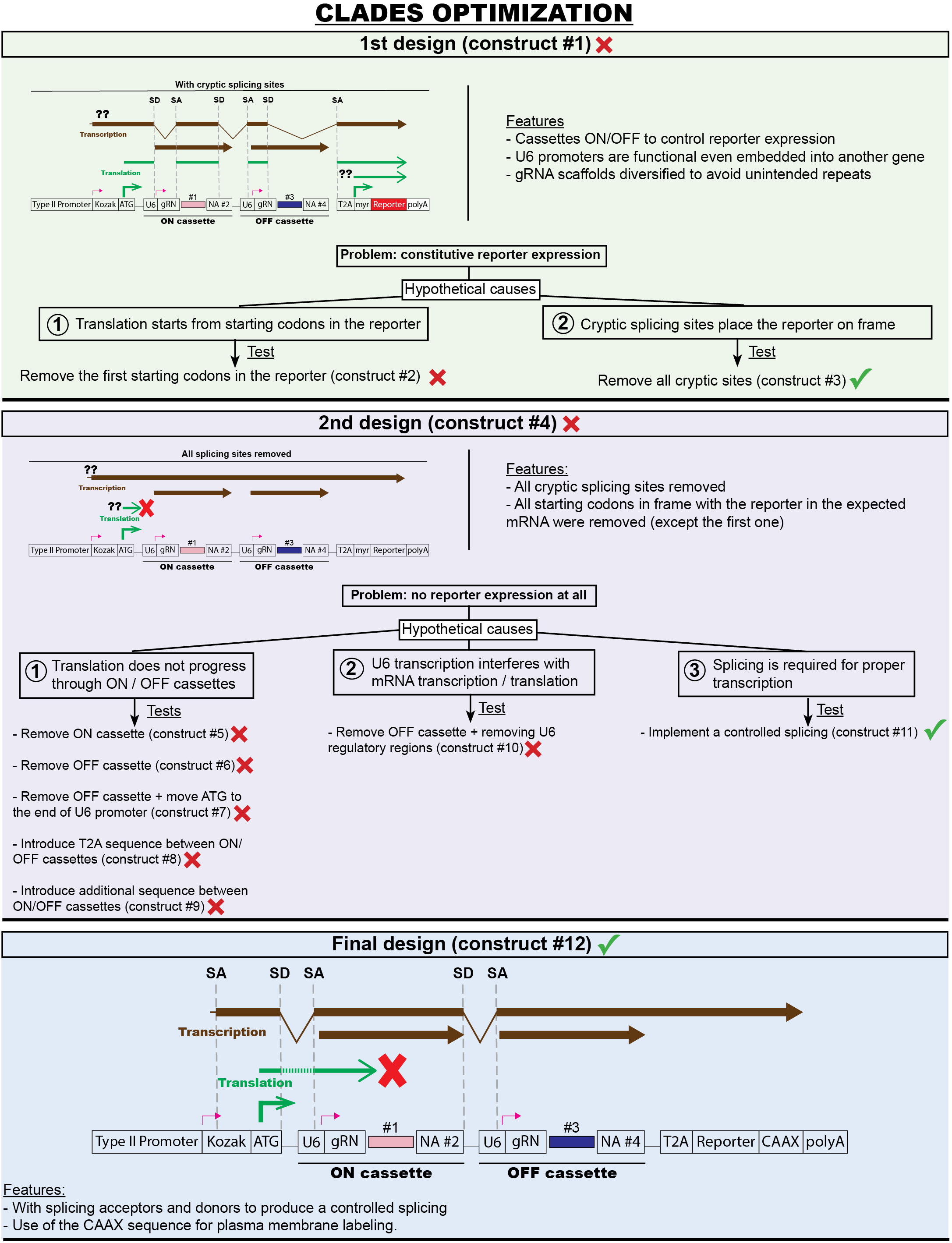
CLADES optimization. Description of the main steps in the generation of a functional CLADES construct. Constructs #1-5, #7 and #11-12 were tested as transgenic flies. Constructs #6, 8, 9-10 were tested in S2 cells. See Supplementary Text and Material and Methods for a detailed description of the optimization process.

**Fig. S6.**
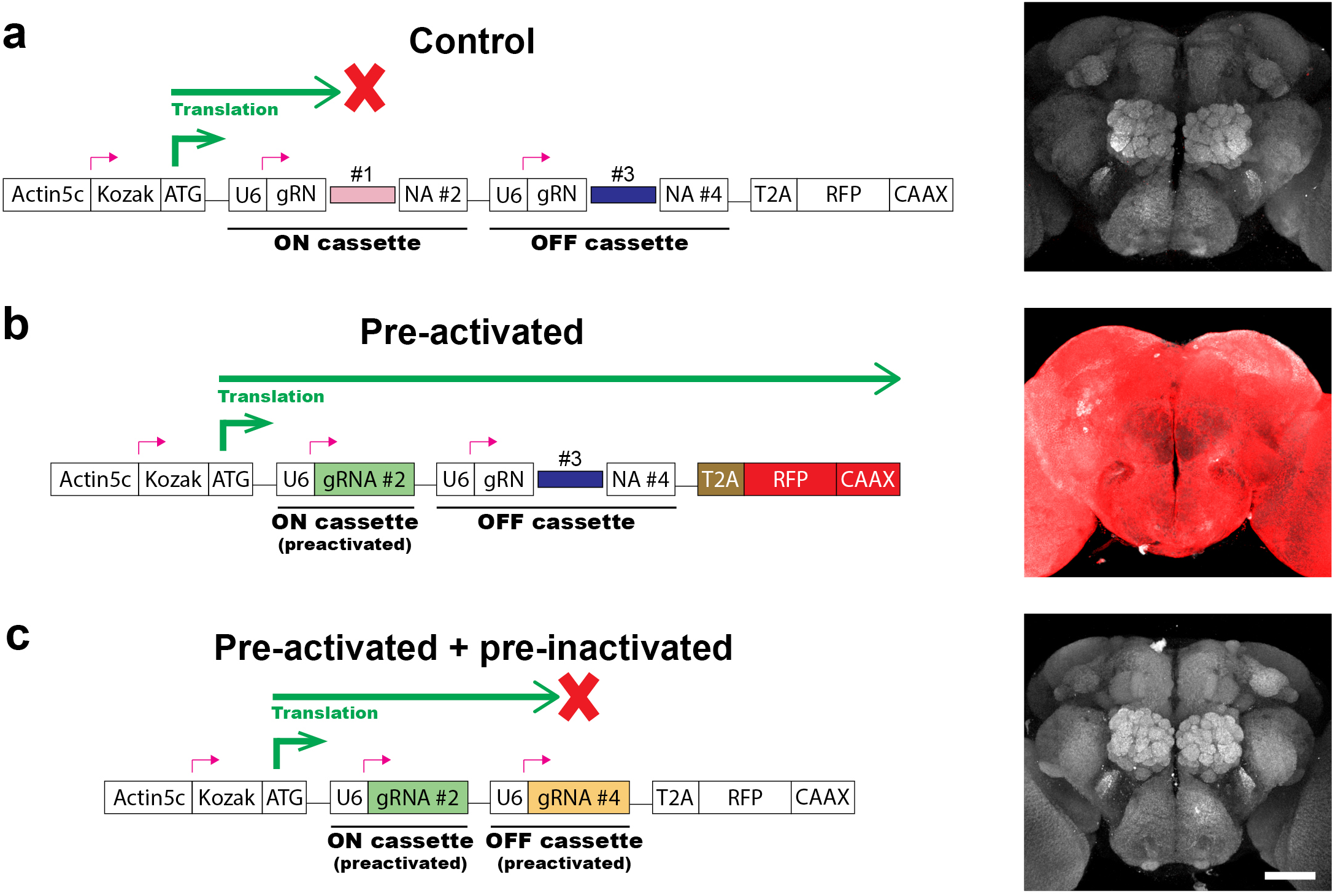
Control constructs showing reporter expression for the different states of CLADES. (A) CLADES construct. In the initial state, no fluorescence is observed as translation stops at the ON cassette. (B) Pre-activated version of CLADES. In this case, the ON cassette sequence is the same as the expected SSA repair outcome. Red fluorescence is ubiquitous since translation progresses to the end of the reporter. (C) Pre-activated+pre-inactivated version of CLADES. Both the sequence for the ON and OFF cassettes is the same as the expected SSA repair outcome. No fluorescence is observed as the translation stops at the OFF cassette. Red and gray, immunohistochemistry for RFP and nc82 respectively. N=12 brains. Scale bar = 50 μm.

**Fig. S7.**
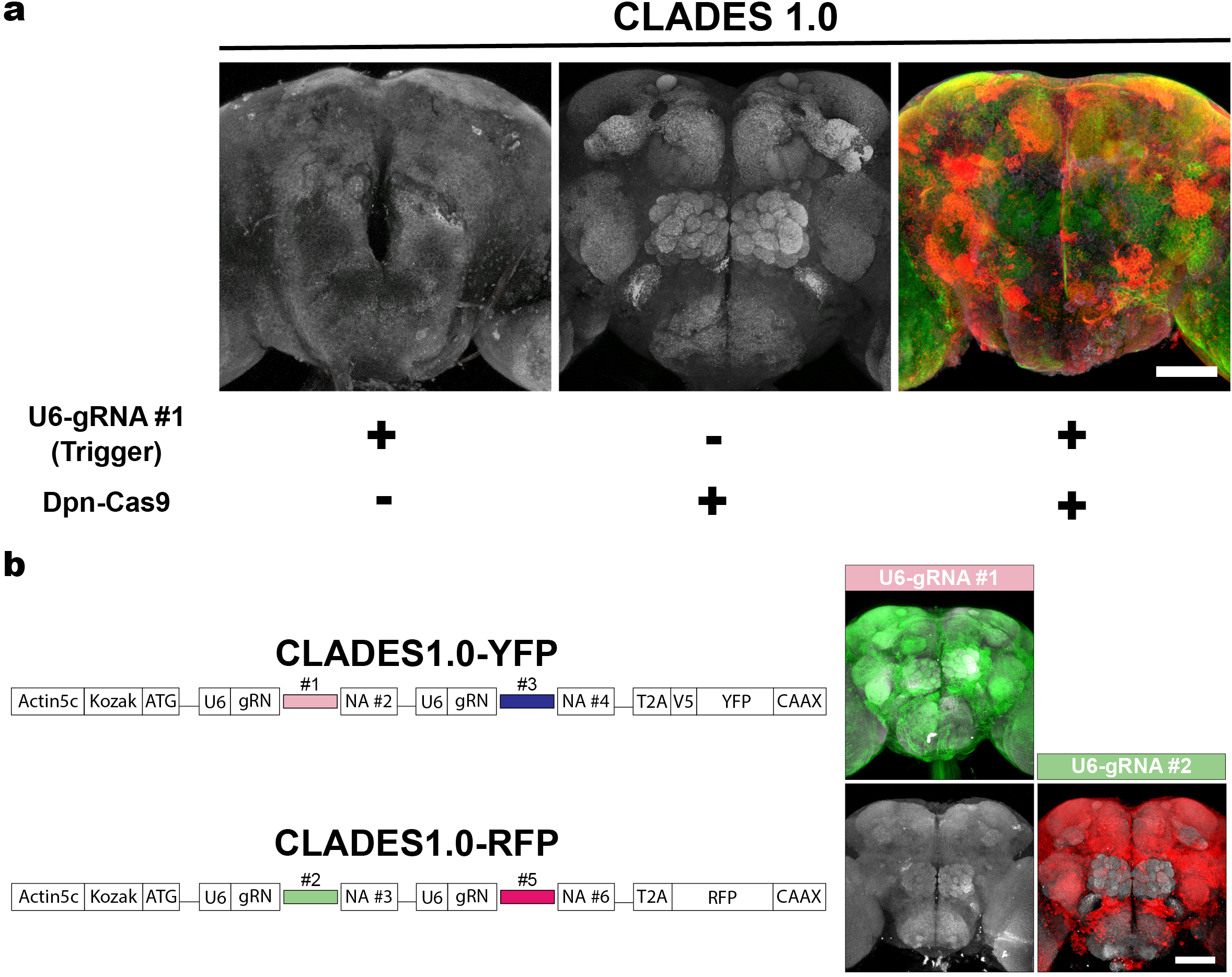
CLADES 1.0 can only be activated by the combination of Cas9 and the trigger gRNA. (A) Triggering CLADES 1.0 requires both the trigger gRNA#1 and Cas9. (B) Only a matching gRNA can trigger each of the CLADES 1.0 reporters. Relevant controls were shown, according to the order in the cascade. Flies bearing Dpn-Cas9 and a U6-gRNA (#1 or #2) were crossed to a fly with CLADES 1.0 (A) or only one of the two CLADES 1.0 constructs (B). Green, red and gray, immunohistochemistry for GFP (YFP), RFP and nc82 respectively. N=24 brains in A and 11 brains for each case in B. Scale bar = 50 μm.

**Fig. S8.**
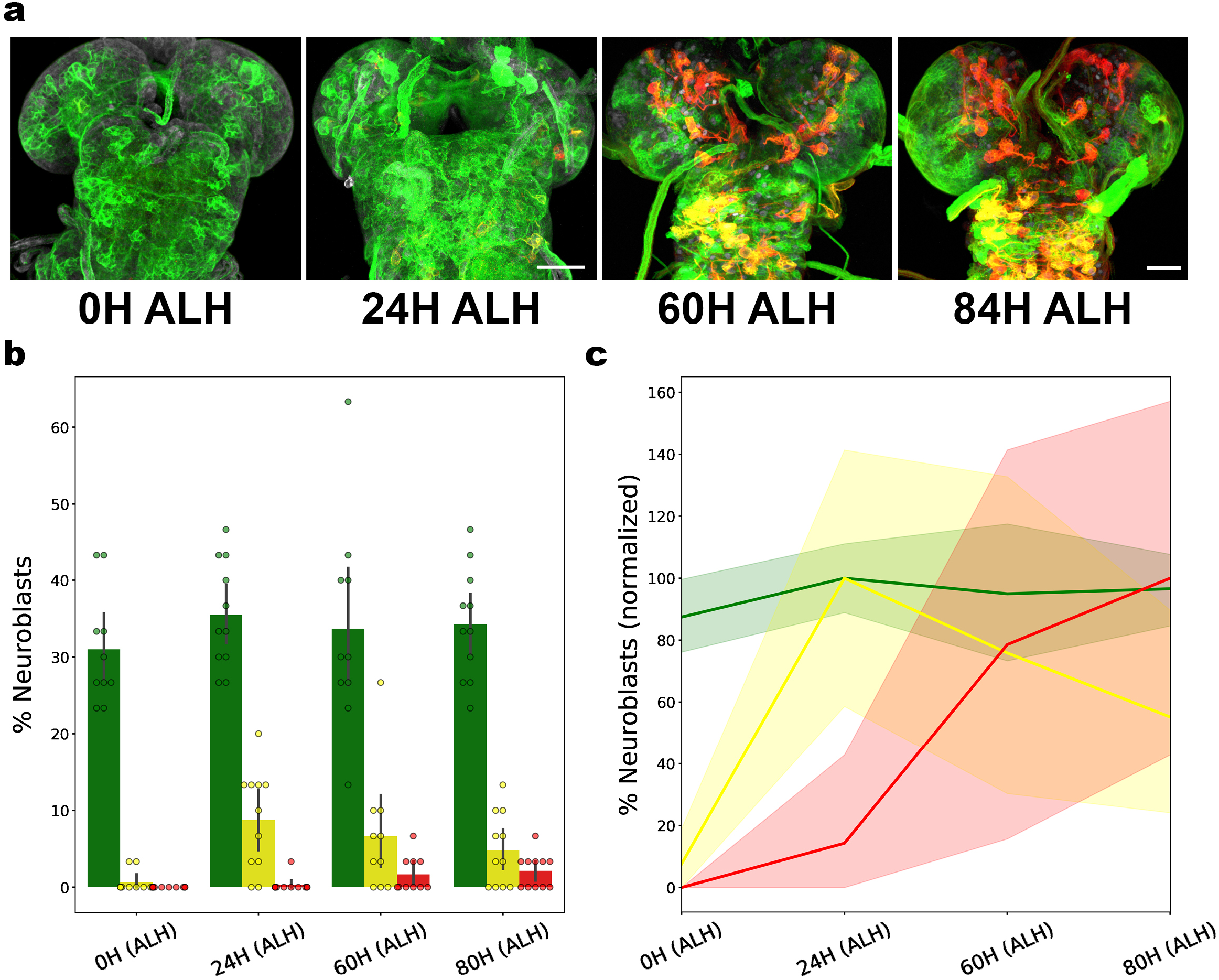
CLADES 1.0 progression over the larval development. (A) Progression of the CLADES 1.0 cascade over the course of larval development, as triggered by the ubiquitous U6-gRNA#1 trigger. Green, red and gray, immunohistochemistry for YFP, RFP and Dpn respectively. (B) Percentage of neuroblasts (n=10 brains, 30 neuroblasts each) exhibiting the different reporter combinations. (C) Normalization of the data shown in (B) to the maximum percentage for each combination of reporters. Error bars and areas around the line plot represent a 95% confidence interval.

**Fig. S9.**
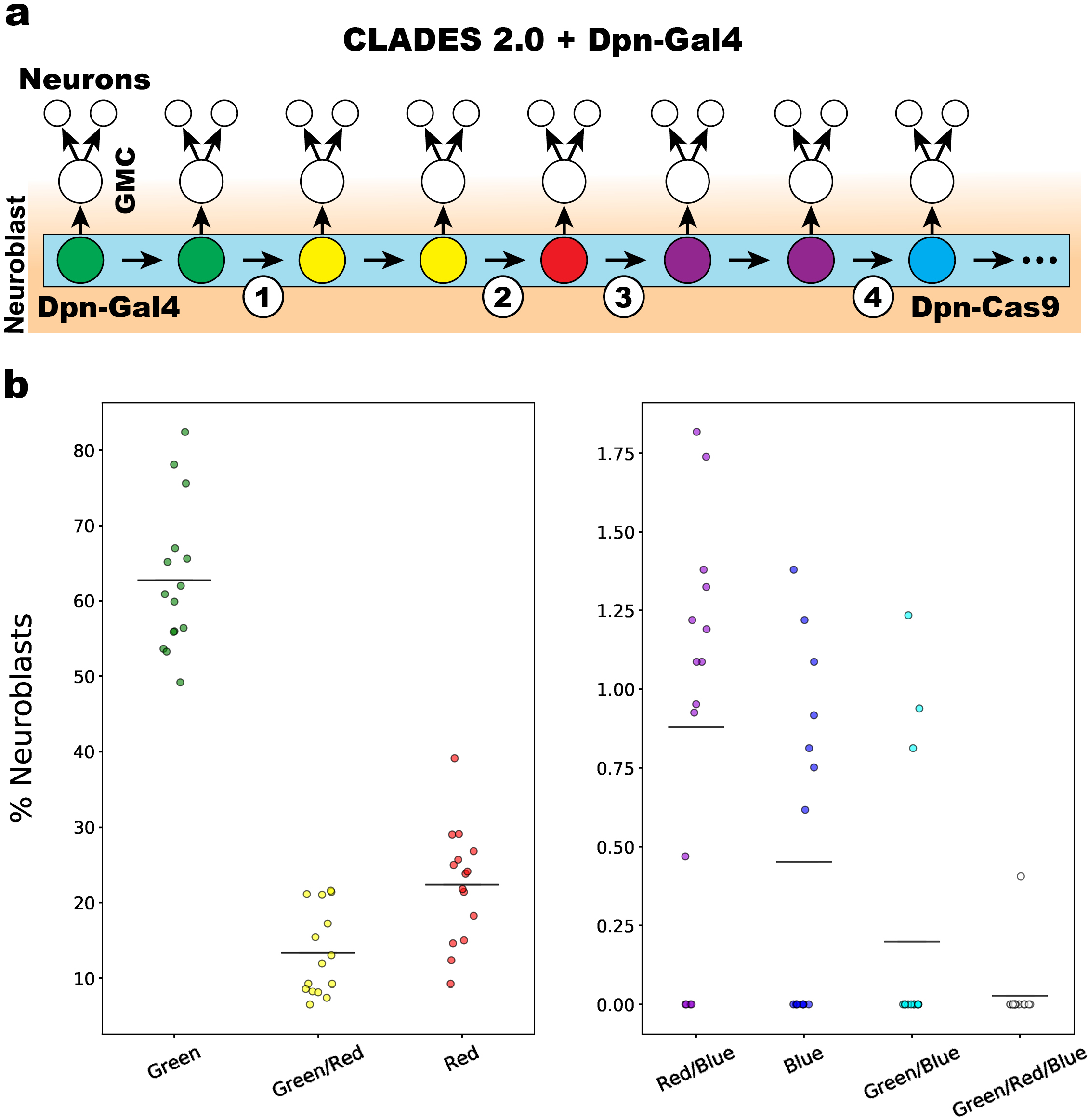
Progression of CLADES 2.0 in neuroblasts. (A) Cartoon illustrating the events occurring after combining CLADES 2.0 and Dpn-GAL4. CLADES progresses in all neuroblasts, driven by Dpn-Cas9. Only those cells expressing Dpn-GAL4 (neuroblasts, with some perdurance in GMC and neurons) are fluorescent. (B) Percentage of neuroblasts expressing each combination of reporters. Horizontal lines represent mean. As the cascade progresses, the proportion of neuroblasts expressing each reporter decays. Only in rare occasions (4 neuroblasts out of ∼3180) unexpected combinations of reporters (blue/green or green/red/blue) were observed, probably due to the minimal leakiness of the conditional gRNAs or the incorrect inactivation of reporters by indels. N=15 brains. Scale bar = 50 μm.

**Table S1.**
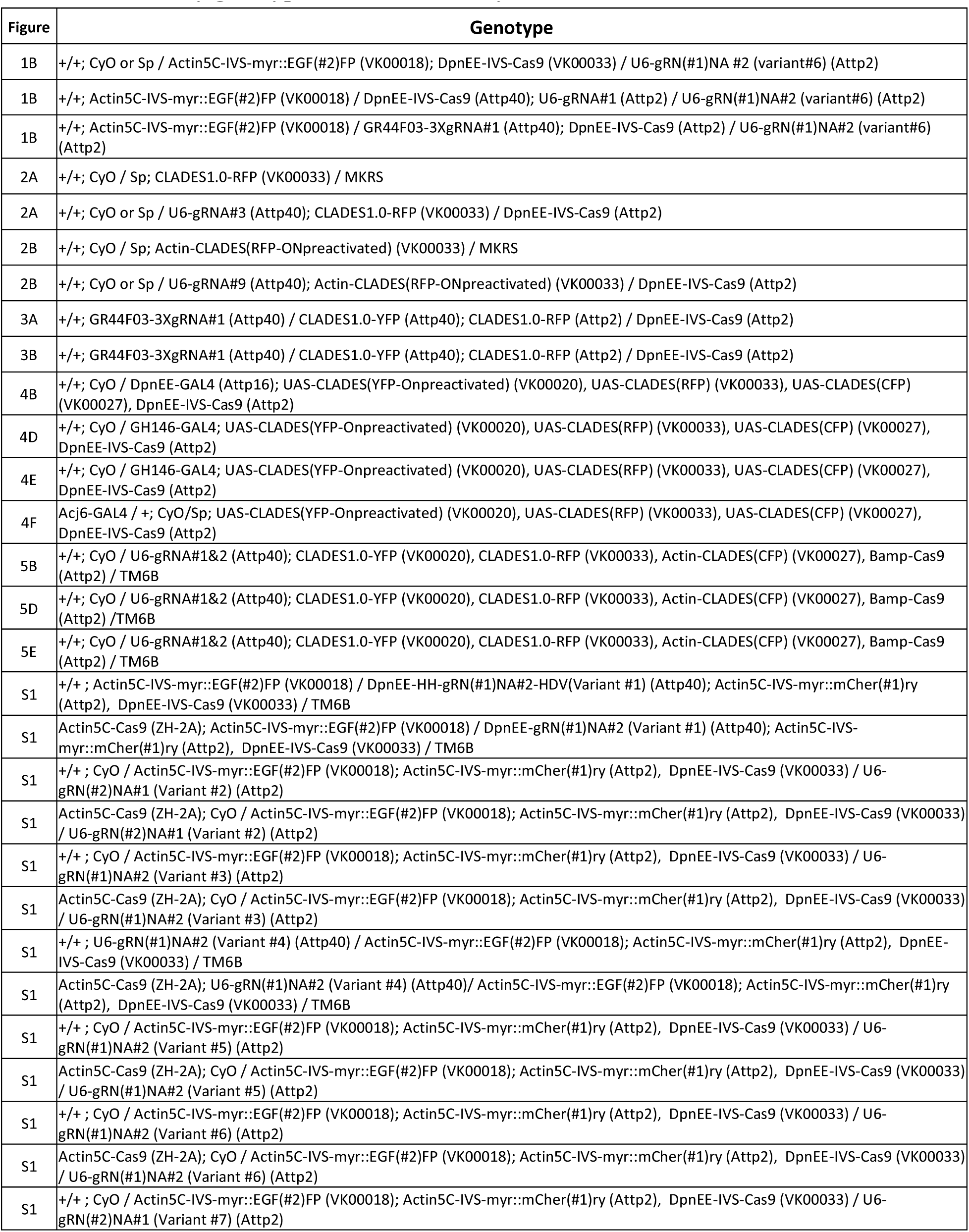
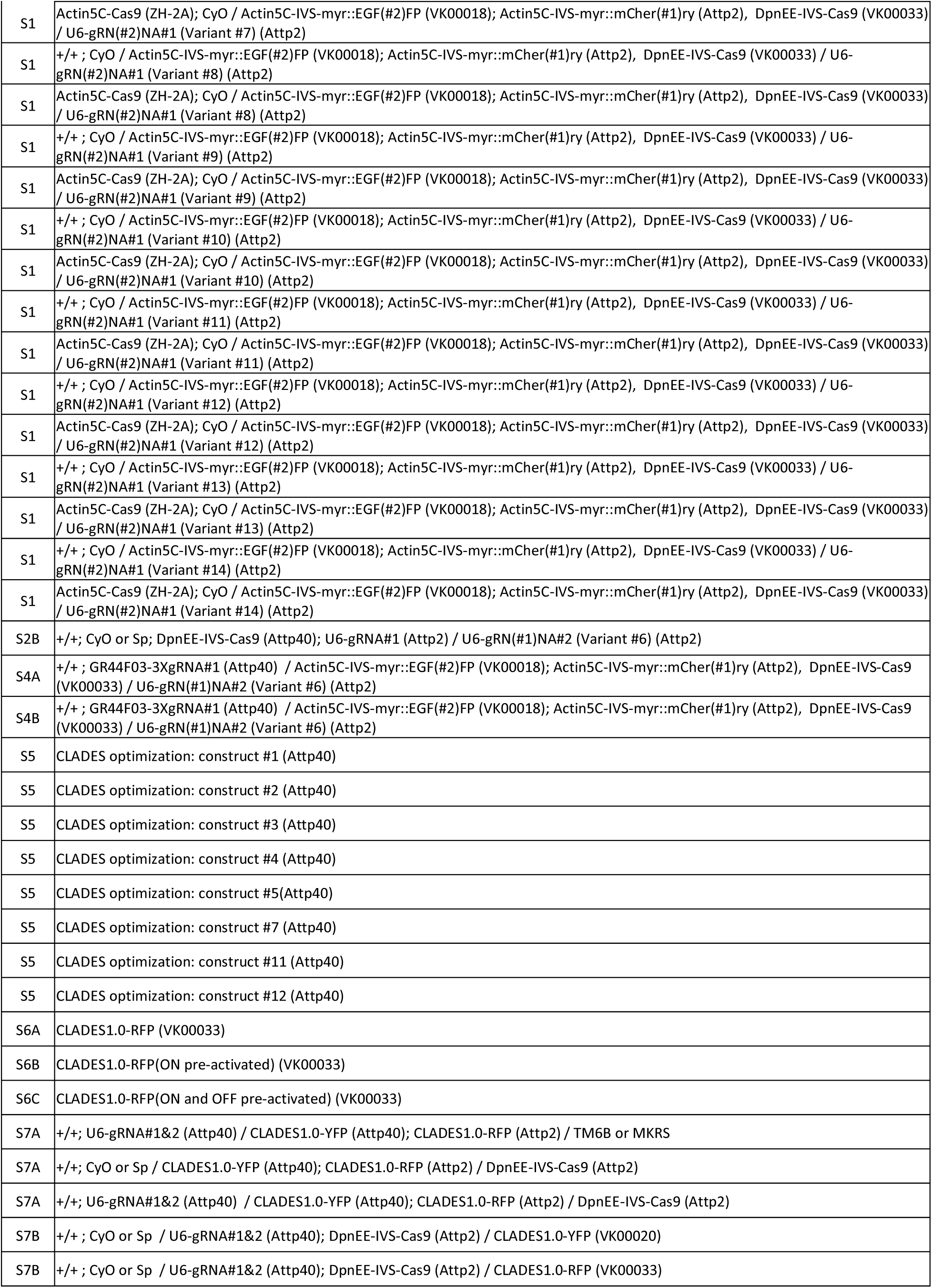
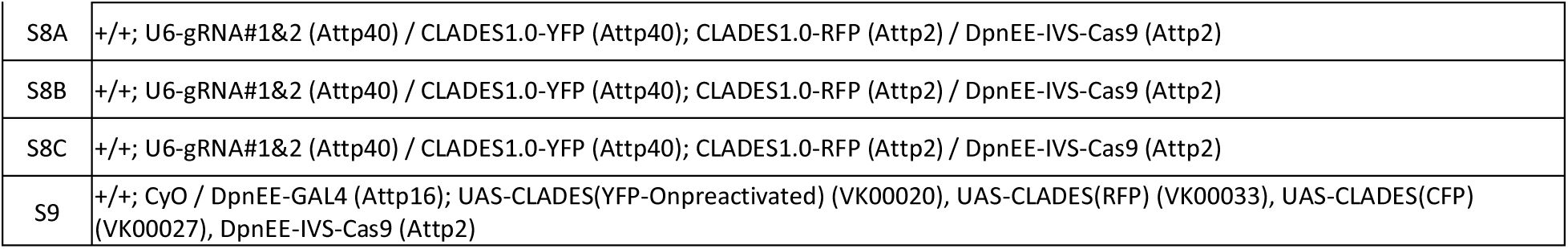
List of fly genotypes used in this study.

**Table S2.**
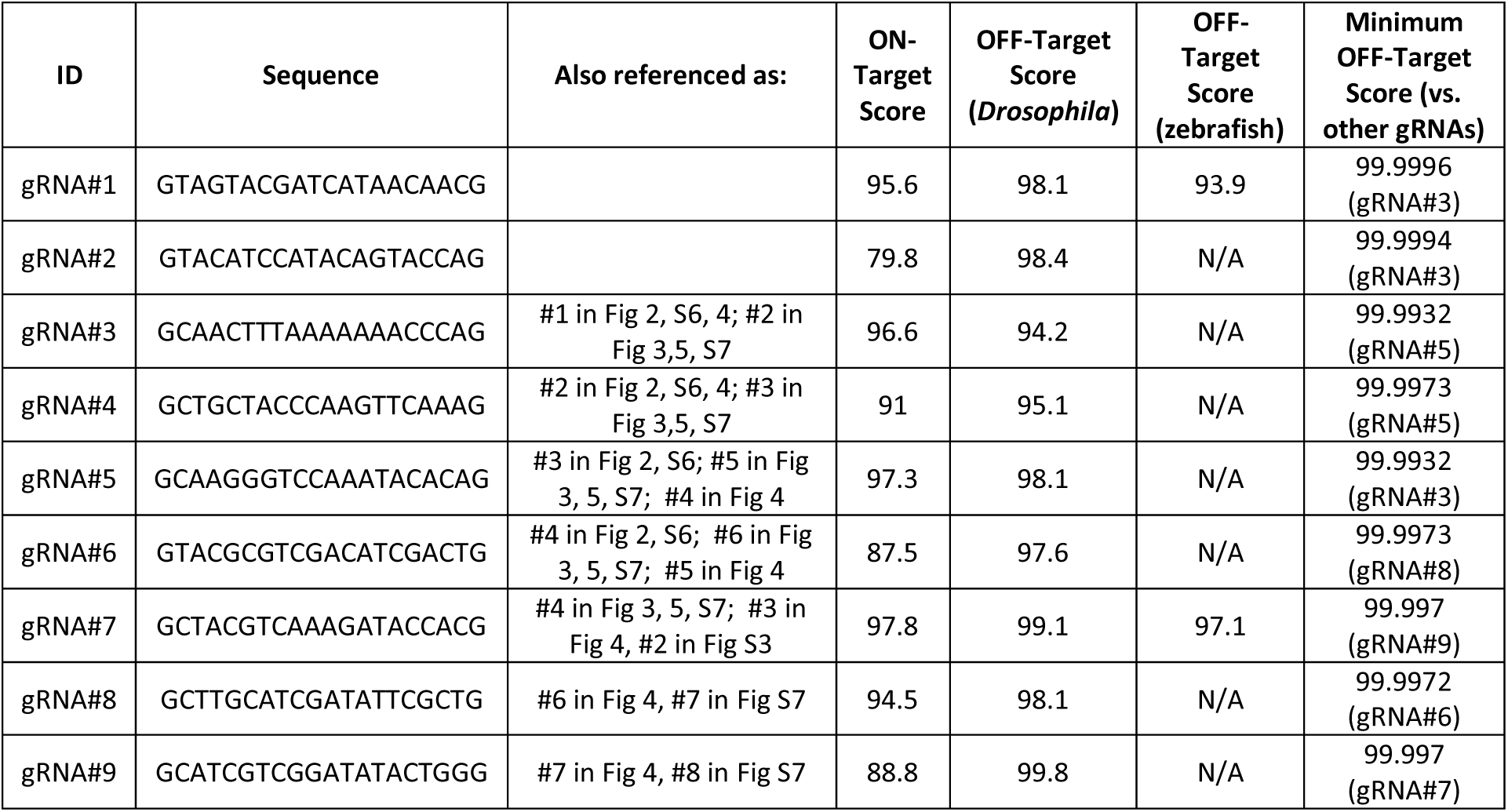
List of gRNAs used in this study. On-Target and OFF-Target scores were calculated as described in Material and Methods.

**Table S3.**
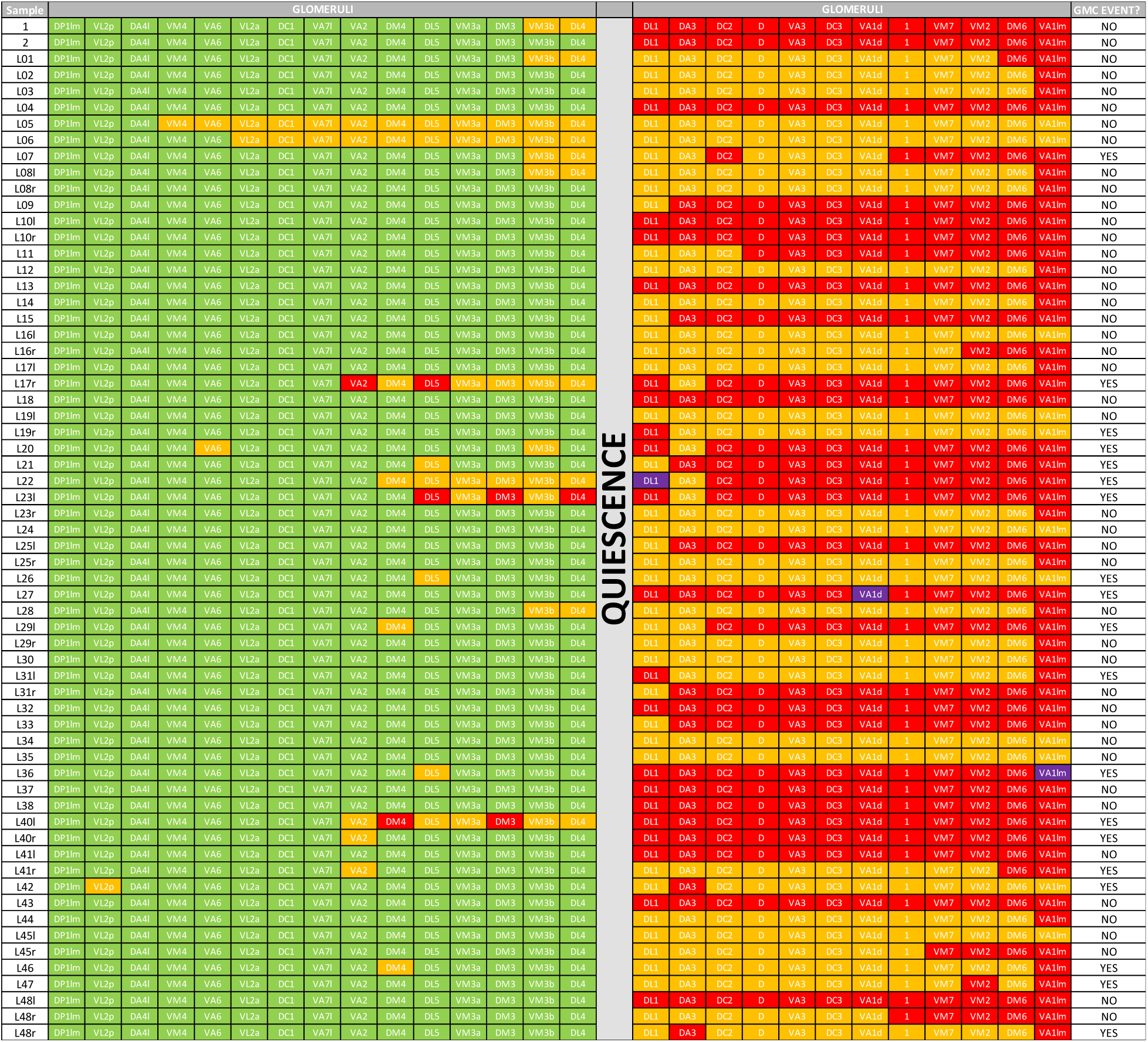
Glomerular projection of Alad1 neurons labeled with GH146-GAL4+Clades 2.0. GMC event refers to Cas9 activity occurring in the GMC rather than in the neuroblast. This leads to progression in the cascade along the neuroblast->GMC->neurons differentiation axis.

